# Vertical gradients in photosynthetic physiology diverge at the latitudinal range extremes of white spruce

**DOI:** 10.1101/2022.05.06.490824

**Authors:** Stephanie C. Schmiege, Kevin L. Griffin, Natalie T. Boelman, Lee A. Vierling, Sarah G. Bruner, Elizabeth Min, Andrew J. Maguire, Johanna Jensen, Jan U. H. Eitel

**Affiliations:** Department of Ecology, Evolution and Environmental Biology, Columbia University, New York, NY 10027, USA; New York Botanical Garden, Bronx, NY 10458; Department of Earth and Environmental Sciences, Columbia University, Palisades, NY 10964, USA; Lamont-Doherty Earth Observatory, Columbia University, Palisades, NY 10964, USA; Department of Natural Resources and Society, College of Natural Resources, University of Idaho, Moscow, ID 83843, USA; McCall Outdoor Science School, College of Natural Resources, University of Idaho, McCall, ID 83638, USA

**Author notes:** CORRESPONDING AUTHOR: Stephanie C. Schmiege, *Email address. Plant Resilience Institute, Michigan State University, East Lansing, MI 48824 USA & Department of Biology, Western University, London, Ontario, Canada N6A 5B7.

**Keywords:** *Picea glauca*, canopy gradients, carbon balance, photosynthesis, photochemical reflectance index, Arctic treeline

## Abstract

Light availability drives vertical canopy gradients in photosynthetic functioning and carbon (C) balance, yet patterns of variability in these gradients remain unclear. We measured light availability, photosynthetic CO_2_ and light response curves, foliar C, nitrogen (N) and pigment concentrations, and the photochemical reflectance index (PRI) on upper and lower canopy needles of white spruce trees (*Picea glauca*) at the species’ northern and southern range extremes. We combined our photosynthetic data with previously published respiratory data to compare and contrast canopy C balance between latitudinal extremes. We found steep canopy gradients in irradiance, photosynthesis, and leaf traits at the southern range limit, but a lack of variation across canopy positions at the northern range limit. Thus, unlike many tree species from tropical to mid-latitude forests, high latitude trees may not require vertical gradients of metabolic activity to optimize photosynthetic C gain. Consequently, accounting for self-shading is less critical for predicting gross primary productivity at northern relative to southern latitudes. Northern trees also had a significantly smaller net positive leaf C balance than southern trees suggesting that, regardless of canopy position, low photosynthetic rates coupled with high respiratory costs may ultimately constrain the northern range limit of this widely distributed boreal species.

**SUMMARY STATEMENT:** Canopy gradients in photosynthetic capacity of white spruce diminish at high compared to low latitudes. Low carbon balance in high latitude trees may determine the extent of northern treeline.

## INTRODUCTION

Illuminating the factors driving species’ distributional ranges, i.e., the boundaries within which a species can grow and reproduce successfully, remains a fundamental ecological, evolutionary, and biogeographic challenge. Extensive research has uncovered a number of factors that play important roles. These include environmental tolerance, niche breadth, metapopulation dynamics, genetic diversity, phenotypic plasticity and dispersal ability (Brown, Stevens & Kaufmnan 1996; Gaston 1996; Lowry & Lester 2006). The diversity of possible factors highlights that no universal cause of species’ distributions has yet emerged, but rather that a combination of historical, physiological, and biotic filters may define a species’ distributional range (Lambers & Oliveira 2019). Nevertheless, the ability of a plant to tolerate the local growth environment remains paramount. This ability is intimately linked to a species’ physiological performance. The physiological processes of photosynthesis and respiration, resulting in the exchange of carbon, are heavily affected by environmental conditions. Together, these two processes determine the carbon balance of a plant which must remain positive if a plant is to grow, survive and reproduce in a given environment. Thus, photosynthesis and respiration fundamentally act as a physiological filter determining a species’ distributional range (Lambers & Oliveira 2019). They are also critical to quantifying carbon exchange as well as the ecological effects of climate change on future carbon sequestration.

The evolution from modeling vegetation as a single “big leaf” to “two big leaves” including distinct parameters for sun and shade leaves, to multi-layer canopies including gradients of physiological function (Field 1983; Sands 1995) has improved estimates of vegetative carbon gain in many ecosystems (Harley & Baldocchi 1995; de Pury & Farquhar 1997; Bonan, Oleson, Fisher, Lasslop & Reichstein 2012; Rogers *et al*. 2017). These multi-layer canopy models are parameterized using physiological and biochemical traits that vary predictably with vertical gradients in environmental conditions such as light, temperature, humidity and vapor pressure deficit throughout the tree crown (Sellers, Berry, Collatz, Field & Hall 1992; Bond, Farnsworth, Coulombe & Winner 1999). Of particular importance to modeling photosynthesis are gradients in light availability. For example, traits such as maximum photosynthesis (*A*_max_) and leaf nitrogen content, as well as leaf mass per area (LMA) have been found to correlate positively with changes in irradiance (Field 1983; Hirose & Werger 1987; Ellsworth & Reich 1993; Niinemets, Kull & Tenhunen 1998; Niinemets 2007) leading to the common finding that high canopy, high light foliage has higher photosynthetic capacity. Pigment pools have similarly been found to vary with canopy position and irradiance with high light foliage frequently containing larger pools of photoprotective pigments that can dissipate excess light energy and protect chlorophyll-filled photosystems (Demmig-Adams & Adams III 1992). Even the ratio of Chl*a*:Chl*b* frequently reflects canopy gradients of light availability because shade-adapted leaves contain more light harvesting antenna complexes (LHCII) associated with photosystem II that are high in Chl*b* (Dale & Causton 1992; Hikosaka & Terashima 1995; Kitajima & Hogan 2003; Ruban 2015; Magney *et al*. 2016; Scartazza, Di Baccio, Bertolotto, Gavrichkova & Matteucci 2016). Thus, the incorporation of canopy gradients into carbon exchange modeling comes from a strong foundation of work that demonstrates how differences in photosynthetic functioning and resource allocation depend on canopy position and light availability.

Other environmental factors may also play a role in determining canopy gradients in photosynthesis. For example, photosynthesis is not only dependent on light availability, but also on water transport to leaves since high photosynthetic rates tend to require high rates of stomatal conductance and transpiration (Wong, Cowan & Farquhar 1978; Schulze, Kelliher, Körner, Lloyd & Leuning 1994). Thus, limitations in water availability may constrain vertical canopy gradients even if light is available in abundance, resulting in shallower canopy gradients in photosynthesis and leaf nitrogen (Peltoniemi, Duursma & Medlyn 2012). Therefore, we cannot assume that light availability is the only driver of canopy gradients without quantifying its effect on photosynthesis, especially in species whose ranges span large latitudinal differences over which environmental conditions can vary dramatically. In this study we examine the strength of the relationships between light availability and canopy gradients of physiological traits across the range extremes of an important boreal species.

White spruce (*Picea glauca* (Moench) Voss) has a massive, transcontinental range distribution stretching from the Forest Tundra Ecotone (FTE) in northern Alaska where it is one of the dominant species, to mixed forests on the eastern seaboard of Canada and New England (Viereck, Van Cleve & Dyrness 1986; Little Jr. 1999; Eitel *et al*. 2019). At the northern edge of the distribution, white spruce are exposed to unique environmental challenges for plant growth and survival including low winter temperatures, shallow soil depth underlain by permafrost, and brief growing seasons. During the growing season, white spruce canopies are exposed to a unique light environment caused by high solar zenith angles and continuous photoperiod (i.e. 24 hours of light for several weeks). Furthermore, tree crown structure at the FTE exhibits an almost ubiquitous tall, vertical, narrow and untapering crown shape (Kuuluvainen 1992) which creates an open canopy forest structure with widely spaced trees. By contrast, white spruce at the southern range extreme are exposed to much lower solar zenith angles and a shorter photoperiod during the growing season. The southern tree crowns are also generally much denser, wider and more tapered, and typically grow with a higher stand density than in the north. These contrasting light environments may impact canopy gradients of light interception, which may, in turn, drive canopy gradients of photosynthetic physiology, biochemistry and pigment concentrations.

In addition to photosynthetic, biochemical and pigment traits, we also examine canopy differences in the photochemical reflectance index (PRI). This index is a more successful photosynthetic proxy than the commonly used normalized difference vegetation index (NDVI) in conifer species because it is sensitive to changes in pigment pools such as the ratio of chlorophylls to carotenoids (Gamon *et al*. 2016). PRI has been successfully employed to track seasonal changes in photosynthetic phenology in coniferous forests (Wong & Gamon 2015a, b; Gamon *et al*. 2016) and at the FTE (Eitel *et al*. 2019, 2020), and a broadband analog Chlorophyll Carotenoid Index (CCI), can be derived from global Moderate Resolution Imaging Spectroradiometer (MODIS) imagery (Gamon *et al*. 2016). Thus, PRI presents a promising prospective avenue for further understanding carbon dynamics across broadly distributed conifer forests.

In this study we focus specifically on the photosynthetic traits of white spruce and their relationship to light environment. However, to gain a full picture of the physiological limits at latitudinally distant range extremes, a carbon balance approach should be employed. In fact, recent work on the physiology of white spruce has focused specifically on the respiratory cost associated with growth and survival at the FTE. In a previous study, we found that white spruce in this environment have extremely high respiration compared to white spruce growing at the southernmost range extreme as well as higher respiration than the average respiration of the boreal biome and needle-leaved evergreen trees (Griffin *et al*. 2021). Griffin et al. (2021) posit that this high respiratory cost may constrain the northern limit of white spruce and the Arctic treeline in general. With the addition of photosynthetic traits measured on the same trees, we are now in a unique position to examine the degree to which this respiratory cost is matched by photosynthetic carbon gains and whether a lower carbon balance may contribute to the northern limits of white spruce’s latitudinal range.

Thus, the overarching aims of this research are twofold: first, to provide a comprehensive assessment of the photosynthetic physiology of white spruce and its relationship to canopy and latitudinal light environment at its range extremes; and second, to examine whether the leaf carbon balance of this species constrains its northernmost range limit. First, we hypothesize that foliar light availability, physiological traits, biochemical traits and PRI will converge across high and low vertical canopy positions in white spruce growing at the FTE. In contrast, steep canopy gradients of light availability and all foliar traits will be apparent at the southernmost range extreme. Second, we hypothesize that the carbon balance of white spruce at the northern location will be much lower than the southern location reflecting the challenging growth environment and providing a physiological mechanism for the northernmost range extreme of this important boreal species.

## MATERIALS AND METHODS

### Site Descriptions

This study was conducted in two locations at the northern and southern range extremes of white spruce (*Picea glauca* (Moench) Voss). In the north, data were collected from six sites chosen to represent the FTE (Eitel *et al*. 2019). These sites are located along a 5.5 km long stretch of the Dalton Highway, in the Brooks Range in northern Alaska (67°59’40.92” N latitude, 149°45’15.84” W longitude). At the FTE, white spruce is the dominant tree species with deciduous shrubs and sedges present in the understory (Eitel *et al*. 2019). The forest structure is relatively open with approximately 407 stems ha^-1^ (Jensen pers. comm.). Three white spruce study trees (with DBH greater than 10cm) were chosen at each site for a total of 18 study trees at the FTE (tree characteristics are described in Table 1). Mean annual precipitation was 485.4 mm and mean annual temperature was -8.1°C as determined from a SNOTEL site at the nearby Atigun Pass (https://wcc.sc.egov.usda.gov/nwcc/site?sitenum=957) between 2007 and 2016. During the Alaska measurement campaign in July 2017, photoperiod ranged from 22 to 24 hours. All measurements were taken at a high and low south-facing canopy position on each of 18 trees for a total of 36 measurements. The low canopy position was at approximately 1.37 m (diameter at breast height), and the high canopy position was approximately 1m below the apical meristem.

**Table 1.** Means ± 1 standard error for characteristics of trees from the Forest Tundra Ecotone (FTE), Alaska (n=18), and Black Rock Forest (BRF), New York (n = 6), including diameter at breast height (DBH; cm), tree height (m), dripline area (m^2^), stems per hectare (means only, as reported in Whitehead et al. (2004) for BRF and from Jensen pers. comm. for the FTE).

| Location | DBH (cm) | Tree Height (m) | Dripline area (m<sup>2</sup>) | Stems ha<sup>-1</sup> |
| --- | --- | --- | --- | --- |
| BRF | 23.10 (1.99) | 9.91 (0.73) | 27.34 (2.00) | 760 |
| FTE | 16.71 (0.94) | 9.32 (0.57) | 3.52 (0.37) | 407 |

At the southern edge of the species’ range, data were collected from six trees located in Black Rock Forest, New York (BRF; 41°24’03.91” N latitude, 74°01’28.49” W longitude; Table 1). Black Rock Forest is an oak-dominated forest with approximately 760 stems ha^-1^ (Whitehead *et al*. 2004). Mean annual precipitation from 1982 – 2010 was 1285 mm and mean annual temperature was 10.9°C (Arguez *et al*. 2012). During our BRF measurement campaign in June 2017, photoperiod was approximately 15 hours. BRF is an oak-dominated northern temperate deciduous forest (Schuster *et al*. 2008; Patterson *et al*. 2018). Measurements were also taken at high and low south-facing canopy positions on each tree. Due to the lower count of available trees, three measurements were taken at the low and high canopy positions on each of the six trees for a total of 36 measurements. As at the FTE, the low canopy position was at approximately 1.37 m and the high canopy position was approximately 1m below the apical meristem.

### Environmental Measurements

Hemispherical photographs were taken with a digital camera (CoolPix 4500, Nikon Corporation, Tokyo Japan) and an attached fisheye lens to assess canopy openness at the high and low canopy positions chosen for each tree (described above). The camera was positioned immediately adjacent to the branch chosen for measurements, with the camera pointing north. Once the camera was level, the hemispherical canopy photo was taken. Using these photographs, light environment was assessed and modeled using the free R code and documentation of ter Steege (2018). These scripts are based on the widely used Hemiphot and Winphot programs (ter Steege 1993, 1997). Modeling of the light environment using this code takes into account GPS location as well as the measurement dates in order to correct for differences in solar angle (see ter Steege (2018) for a fuller description). Average Photosynthetic Photon Flux Density (PPFD ; mol m^-2^ day^-1^) at the high and low canopy positions was calculated by averaging the diurnal light courses over the measurement campaign days (June 21 – June 27 at BRF, and July 4 – July 20 at the FTE). At the FTE, air temperature (°C) was collected using a meteorological sensor (VP-4, METER, Pullman, WA) and local solar open sky radiation readings were collected using a PYR Solar Radiation Sensor (METER, Pullman, WA). Solar radiation (W m^-2^) was converted to PPFD (μmol m^-2^ s^-1^) by multiplying by 2.02 (Mavi & Tupper 2004). At BRF, air temperature and local open sky PPFD were collected from a nearby BRF-run weather station (https://blackrock.ccnmtl.columbia.edu/portal/weather/) equipped with a temperature/humidity sensor (HC2S3, Rotronic, Hauppauge, NY) and a quantum sensor (LI-190SB Quantum Sensor, LICOR, Lincoln, NE).

### Gas Exchange Measurements

Gas exchange was measured at high and low canopy positions on each tree at the two locations. At both locations, measurements were taken on a branch tip inserted into the cuvette. At BRF, all measurements were taken on the tree with high canopy reached using an articulating boom lift (600AJ, JLG, McConnellsburg, PA). At the FTE, low canopy branches were measured on the tree, but high canopy branches had to be detached using a pole-clippers (Fiskars fiberglass pole pruner, Fiskars Brands Inc, Middleton, WI) as there was no way to reach samples otherwise. Detaching foliage for physiological measurements is common, with no observable differences usually found between attached and detached samples (Akalusi, Meng & Bourque 2021). For detached samples, cut branches at least 30 cm in length were placed immediately in water, and recut underwater. All measurements on cut branches were taken within 12 hours. CO_2_ (*A-C*_i_) and light response (*A-Q*) curves were collected using two portable photosynthesis systems (LI-6800, LiCor, Lincoln, NE). Needles were sealed in a cuvette in which the temperature and relative humidity were set to 20°C and 50% humidity at the FTE, and 25°C, and 60% humidity at BRF in order to mirror ambient conditions in each location. For the *A-C*_i_ curves, irradiance was kept constant at 1000 PPFD and the CO_2_ concentrations were stepped through the following concentrations (ppm): 1200, 1000, 800, 600, 400, 300, 200, 100, 75, 50, 25, 0. Values of *V*_cmax_ (the maximum rate of RuBisCO carboxylation) and *J*_max_ (the maximum rate of electron transport) were determined according to the photosynthesis model of Farquhar, von Caemmerer and Berry (Farquhar, von Caemmerer & Berry 1980) and adjusted to 25°C using the plantecophys package in R (Duursma 2015).

For the *A-Q* curves, reference CO_2_ concentration was set to 400 ppm and photosynthesis was measured at the light intensities: 1800, 1500, 1200, 1000, 800, 400, 200, 150, 100, 90, 80, 75, 70, 65, 60, 55, 50, 45, 40, 35, 30, 25, 20, 15, 10, 5, 0 μmol m^-2^ s^-1^. After 15 minutes at zero irradiance, respiration in the dark (*R*_D_) was recorded. Maximum likelihood estimation (executed in the bblme package for R (Bolker & R Development Core Team 2020)) was used to fit the following model (describing a non-rectangular hyperbola) to the data (Herrmann, Schwartz & Johnson 2020):

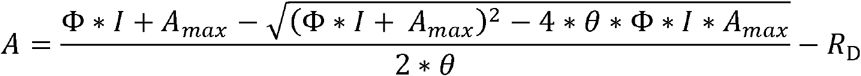

Where *A* is the measured rate of photosynthesis, is the curvature factor and *I* is the irradiance. Parameters extracted and calculated from the fit of the model to each light curve included the maximum rate of CO_2_ fixation (*A*_max_), photosynthesis at 1500 μmol m^-2^ s^-1^ (*A*_1500_), the apparent quantum yield (Φ), the light compensation point (LCP – the light level required for zero net carbon flux) and the light saturation point (LSP – the light level required to obtain 75% of *A*_1500_).

### Respiration in the Light

Respiration in the light (*R*_L_) was obtained from the *A*-*Q* curves according to the methods of Kok (1948); Sharp, Matthews & Boyer (1984); and Heskel, Atkin, Turnbull & Griffin (2013). These curves contain a subtle change in slope near the light compensation point (the light intensity where photosynthesis and respiration equal each other). The break point where this change in slope occurs divides the photosynthetic response to light into two linear sections. Extrapolating from the upper linear portion of the curve to the y-axis highlights an alternate respiration value which was interpreted by Bessel Kok to be the amount of respiration occurring in the light (Kok 1948; Heskel *et al*. 2013; Tcherkez *et al*. 2017).

An important assumption of the Kok method is that carbon assimilation is only influenced by light. Consequently, corrections were made to account for any changes in internal CO_2_ concentration (*C*_i_) following the methods of Kirschbaum & Farquhar (1987) (see Ayub *et al*. (2011) for a full description). It is possible that *R*_L_ measured with the Kok method may be influenced by factors such as mesophyll conductance (Farquhar & Busch, 2017; *n*.*b*. Buckley *et al*., 2017); however, such investigations are beyond the scope of this study.

### Leaf Trait Measurements

After collecting *A*-*C*_i_ and *A-Q* curves, needles were detached from the branch, photographed to determine projected surface area (cm^2^) using ImageJ (Schnieder, Rasband & Eliceiri 2012), placed into coin envelopes and oven dried at 60°C for 48 hours. Leaves were reweighed post-drying for leaf dry mass (g). Specific leaf area (SLA; cm^2^g^-1^) was calculated. Leaf samples were then ground using a ball mill (SPEX 8000 Mixer/Mill, Metuchen, NJ, USA) and prepared for analysis of % carbon and % nitrogen using a carbon-nitrogen flash analyzer (CE Elantech, Lakewood, NJ, USA). Using these percentages, foliar nitrogen per area (N_area_; mg cm^-2^) and the carbon to nitrogen ratio (C:N) were calculated.

### Pigment measurements

Needles growing on branches adjacent to those selected for gas exchange at both high and low canopy positions were selected for pigment analysis. Branches were wrapped in wet paper towels and foil, and transported back to the laboratory in a cooler with ice. Foliar pigment extraction began no later than three hours after returning from the field, and any samples not processed in this time were frozen until pigment extraction could take place.

Between 5 and 10 needles were removed from each branch and their wet weight and needle area measured. Needles were ground in 100% acetone until no identifiable green needle tissue remained. A small amount of sand, MgCO_3_ and ascorbic acid were added to help with grinding and to prevent acidification and pigment degradation. The extracts were then centrifuged and diluted. Using a spectrophotometer (Go Direct SpectroVis Plus Spectrophotometer, Vernier, OR, USA), the absorbance of the supernatant was measured at 470, 645, 662 and 710 nm. Pigment pools were calculated on a leaf area basis including chlorophyll *a* (Chl*a*_area_), chlorophyll *b* (Chl*b*_area_), and bulk carotenoids (Car_area_; ug/cm^2^ for all pools) were calculated according to Lichtenthaler (1987). The total chlorophyll pool size (Chl_area_ = Chl*a*_area_ + Chl*b*_area_) and ratios Chl*a*:Chl*b*_area_ and Chl:Car_area_ were also calculated.

### Spectral reflectance: PRI

Needle spectral reflectance was measured with a spectroradiometer (UniSpec SC, PP Systems, Haverhill MA, USA) with an attached fiber optic probe (UNI400, PP Systems, Haverhill MA, USA) and specialized needle clip (UNI501, PP Systems, Haverhill MA, USA). The spectroradiometer allowed sampling of the spectrum between 310 and 1100 nm with a 3.3 nm sampling interval which was afterwards reduced to 1 nm using linear interpolation. After collecting dark and white standard (Spectralon, LabSphere, North Sutton, NY, USA) reference measurements, three spectral measurements were made on each branch to capture spatial heterogeneity. Reflectance values were calculated from the spectral measurements by dividing each foliage measurement by the measurement from the white standard. PRI was calculated according to the equation: R531 R570 / R531 R570 (Gamon, Penuelas & Field 1992; Penuelas, Filella & Gamon 1995) where R531 and R570 are the reflectances at 531 nm and 570 nm, respectively.

### Statistical analyses

Relationships between measured foliar parameters (physiology, pigments and leaf traits), and integrated daily PPFD and location (BRF or the FTE) were assessed using linear mixed effects models. The interaction between PPFD and location was included so as to examine the variations in slope between locations. Individual tree (FTE n = 18, BRF n = 6) was included as a random effect to account for repeat measures on trees (Parameter ∼ PPFD * location + (1|tree)). If necessary, parameters were log-transformed to meet assumptions of normality of the residuals. Parameter estimates were considered significantly different from zero if P<0.05. Linear mixed effects analysis was conducted using the lme4 package (Bates, Maechler, Bolker & Walker 2015) with the lmerTest package loaded to provide p-values (Kuznetsova, Brockhoff & Christensen 2017). All analyses took place in R v. 4.1.0 (R Core Team 2022).

The effects of canopy position and location on foliar traits were also assessed using mixed effects models implemented in the lme4 package (Bates *et al*. 2015). Again, individual tree was included as a random effect to account for repeat measures on trees. Pairwise comparisons between canopy position (high or low) and location (FTE or BRF) were assessed using the emmeans package in R and found to be significant if P<0.05 (Lenth 2020). All data reported below are means ± SE. Finally, the relationships between physiological parameters and either N_area_ or total chlorophyll (Chl+Car_area_) were also examined using linear mixed effects models. A description of the full models can be found in the Supplemental Information Methods S1.

## RESULTS

### Tree Characteristics and Growth Environment

Tree characteristics for the trees measured in this study have been previously reported in Griffin *et al*. (2022) but have been summarized in Table 1. Briefly, trees from BRF tended to have a larger diameter at breast height (DBH; measured at 1.37 m from the ground) with BRF trees having an average diameter of 23.1±1.99 cm and FTE trees having an average diameter of 16.7±0.94 cm (Table 1). Trees from the two locations were also extremely different in their average dripline areas (27.34±2.00 m^2^ at BRF vs. 3.52±0.37 m^2^ at the FTE; Table 1). Nevertheless, despite these differences trees from the two locations had very similar average heights (9.91±0.73 m at BRF and 9.32±0.57 m at the FTE; Table 1).

Irradiance at the range extremes of white spruce, from the northernmost location at the FTE in Alaska, to the southernmost location at BRF in New York, showed pronounced differences in both intensity and day length (Fig 1a, b, d). Average ambient PPFD over the measurement campaigns (Fig 1a; orange) estimated from daily diurnal light courses modeled from canopy photos (e.g., July 4^th^ 2017; Fig 1b) was 36% higher at BRF than at the FTE (54.4±0.03 compared to 39.9±0.06 mol m^-2^ day^-1^). However, day length was much longer at the FTE than at BRF (Fig 1b). At the canopy positions where physiological and biochemical measurements were made, average daily PPFD was significantly greater at high canopy positions than low canopy positions at both BRF and the FTE (34.97±0.16 vs. 9.20±0.13 mol m^-2^ day^-1^ at BRF, and 29.24±0.16 vs. 21.14±0.16 mol m^-2^ day^-1^ at the FTE; Fig 1a). Even so, the difference in the daily mean PPFD experienced from the upper to the lower canopies was much smaller at the FTE than at BRF (only 8 mol m^-2^ day^-1^ at the FTE vs. 25 mol m^-2^ day^-1^ at BRF). This narrowing of the irradiance gradient across vertical canopy positions at the FTE was decoupled from canopy height, given that trees at both locations were of similar heights.

**Figure 1.**
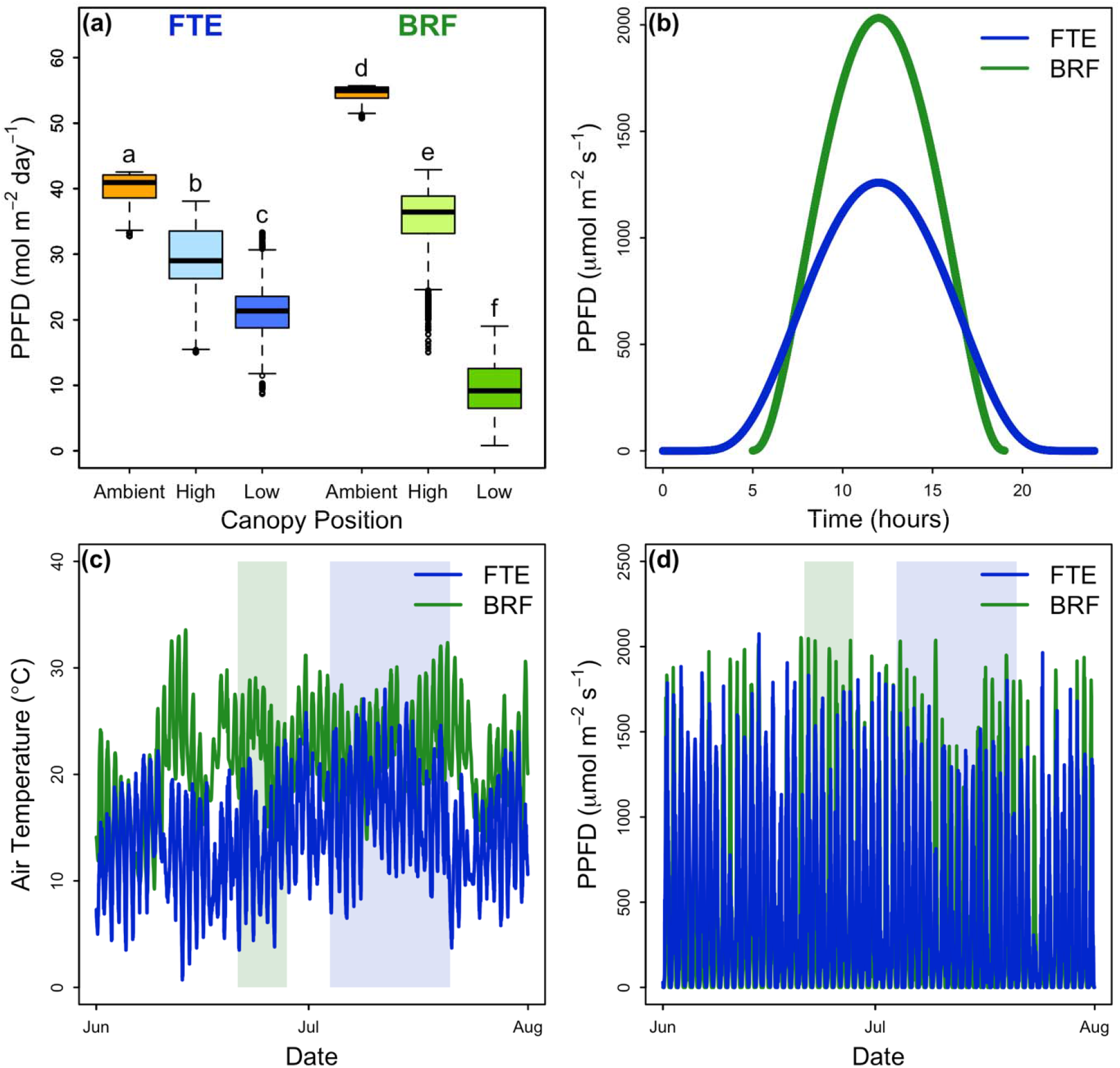
(a) Average daily PPFD calculated from canopy photos over the study period (June-July) at the Forest Tundra Ecotone (FTE), Alaska, and Black Rock Forest (BRF), New York. Ambient (above canopy) PPFD is shown in orange. Boxplots show the median and first and third quartiles. Whiskers display the range of groups with individual points representing outliers falling outside 1.5 times the interquartile range. Different letters represent significant differences between locations and canopy positions (P<0.05). (b) Ambient PPFD projected from canopy photos during one day (July 4, 2017) at both locations. (c) Air temperature (°C) measured at BRF and the FTE for June and July 2017. (d) Ambient PPFD measured from field instruments at BRF and the FTE for June and July 2017. Green and blue shaded areas in 1c and 1d denote the dates of the study campaigns at BRF and the FTE, respectively.

In addition to differences in the light environment, there were also pronounced differences in the daytime air temperatures between the two locations (Fig 1c). During the respective study campaigns (as marked by the green (BRF) and blue (FTE) shaded areas (Fig 1c), BRF mean daytime temperature was 23.4°C, with a minimum temperature of 12.8°C and a maximum temperature of 29.1°C, whereas the FTE mean temperature was 17°C, with a minimum temperature of 6.5°C and a maximum temperature of 28°C. Thus, the FTE experienced colder temperatures and greater temperature variability.

### Relationships between light environment and gas exchange, biochemistry, pigments and PRI

In general, significant relationships were found between average PPFD and physiological traits, biochemical traits, pigment concentrations and PRI at BRF. In contrast, few significant relationships were found between average PPFD and foliar traits at the FTE (Table 2).

**Table 2.**
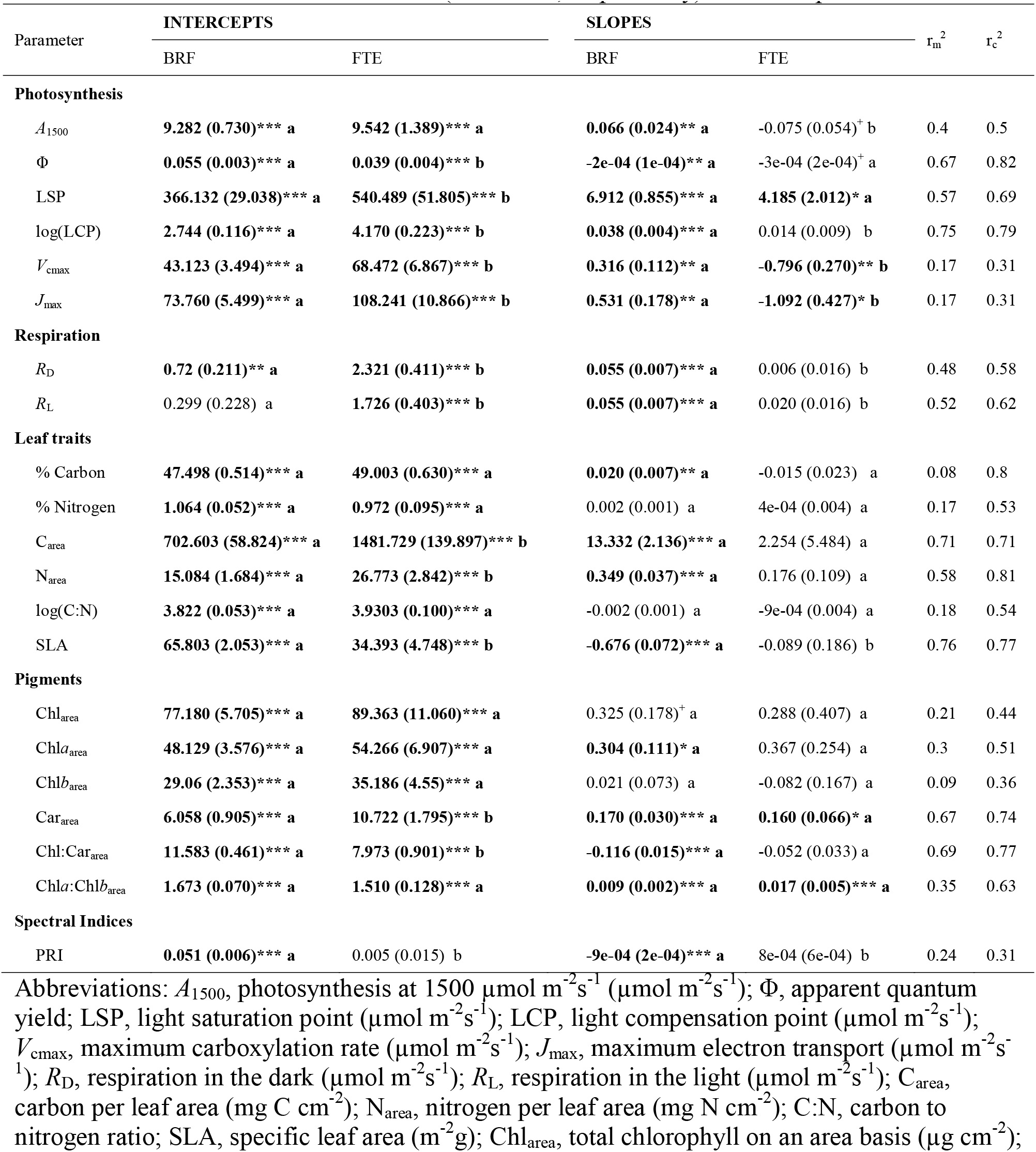
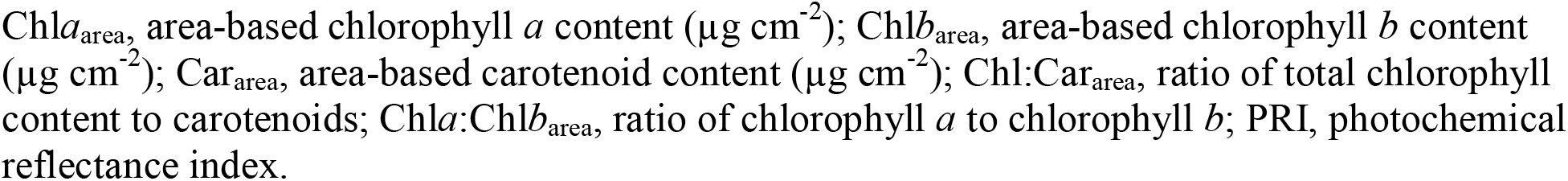
Parameter estimates for the linear mixed effects models in which each physiological, leaf and spectral trait was modeled as a function of integrated average daily photochemical photon flux density (PPFD) and location (either Black Rock Forest (BRF), New York, or the Forest Tundra Ecotone (FTE), Alaska) with a random effect for each individual tree (Parameter ∼ PPFD * location + (1|tree)). Values in parentheses are SEs. Bolded and starred values are those that are significantly different from zero (***P<0.001; **P<0.01; *P<0.05; ^+^P<0.1). Significant differences between locations are indicated by different letters (P<0.05). Marginal and conditional coefficients of determination (r_m_^2^ and r_m_^2^, respectively) are also reported.

| Parameter | INTERCEPTS | | SLOPES | | $r_m^2$ | $r_c^2$ |
| --- | --- | --- | --- | --- | --- | --- |
|  | BRF | FTE | BRF | FTE |  |  |
| Photosynthesis |  |  |  |  |  |  |
| $A_{1500}$ | 9.282 (0.730)*** a | 9.542 (1.389)*** a | 0.066 (0.024)** a | -0.075 (0.054)<sup>+</sup> b | 0.4 | 0.5 |
| $\Phi$ | 0.055 (0.003)*** a | 0.039 (0.004)*** b | -2e-04 (1e-04)** a | -3e-04 (2e-04)<sup>+</sup> a | 0.67 | 0.82 |
| LSP | 366.132 (29.038)*** a | 540.489 (51.805)*** b | 6.912 (0.855)*** a | 4.185 (2.012)* a | 0.57 | 0.69 |
| log(LCP) | 2.744 (0.116)*** a | 4.170 (0.223)*** b | 0.038 (0.004)*** a | 0.014 (0.009) b | 0.75 | 0.79 |
| $V_{cmax}$ | 43.123 (3.494)*** a | 68.472 (6.867)*** b | 0.316 (0.112)** a | -0.796 (0.270)** b | 0.17 | 0.31 |
| $J_{max}$ | 73.760 (5.499)*** a | 108.241 (10.866)*** b | 0.531 (0.178)** a | -1.092 (0.427)* b | 0.17 | 0.31 |
| Respiration |  |  |  |  |  |  |
| $R_D$ | 0.72 (0.211)** a | 2.321 (0.411)*** b | 0.055 (0.007)*** a | 0.006 (0.016) b | 0.48 | 0.58 |
| $R_L$ | 0.299 (0.228) a | 1.726 (0.403)*** b | 0.055 (0.007)*** a | 0.020 (0.016) b | 0.52 | 0.62 |
| Leaf traits |  |  |  |  |  |  |
| % Carbon | 47.498 (0.514)*** a | 49.003 (0.630)*** a | 0.020 (0.007)** a | -0.015 (0.023) a | 0.08 | 0.8 |
| % Nitrogen | 1.064 (0.052)*** a | 0.972 (0.095)*** a | 0.002 (0.001) a | 4e-04 (0.004) a | 0.17 | 0.53 |
| $C_{area}$ | 702.603 (58.824)*** a | 1481.729 (139.897)*** b | 13.332 (2.136)*** a | 2.254 (5.484) a | 0.71 | 0.71 |
| $N_{area}$ | 15.084 (1.684)*** a | 26.773 (2.842)*** b | 0.349 (0.037)*** a | 0.176 (0.109) a | 0.58 | 0.81 |
| log(C:N) | 3.822 (0.053)*** a | 3.9303 (0.100)*** a | -0.002 (0.001) a | -9e-04 (0.004) a | 0.18 | 0.54 |
| SLA | 65.803 (2.053)*** a | 34.393 (4.748)*** b | -0.676 (0.072)*** a | -0.089 (0.186) b | 0.76 | 0.77 |
| Pigments |  |  |  |  |  |  |
| $Chl_{area}$ | 77.180 (5.705)*** a | 89.363 (11.060)*** a | 0.325 (0.178)<sup>+</sup> a | 0.288 (0.407) a | 0.21 | 0.44 |
| $Chla_{area}$ | 48.129 (3.576)*** a | 54.266 (6.907)*** a | 0.304 (0.111)* a | 0.367 (0.254) a | 0.3 | 0.51 |
| $Chlb_{area}$ | 29.06 (2.353)*** a | 35.186 (4.55)*** a | 0.021 (0.073) a | -0.082 (0.167) a | 0.09 | 0.36 |
| $Car_{area}$ | 6.058 (0.905)*** a | 10.722 (1.795)*** b | 0.170 (0.030)*** a | 0.160 (0.066)* a | 0.67 | 0.74 |
| $Chl:Car_{area}$ | 11.583 (0.461)*** a | 7.973 (0.901)*** b | -0.116 (0.015)*** a | -0.052 (0.033) a | 0.69 | 0.77 |
| $Chla:Chlb_{area}$ | 1.673 (0.070)*** a | 1.510 (0.128)*** a | 0.009 (0.002)*** a | 0.017 (0.005)*** a | 0.35 | 0.63 |
| Spectral Indices |  |  |  |  |  |  |
| PRI | 0.051 (0.006)*** a | 0.005 (0.015) b | -9e-04 (2e-04)*** a | 8e-04 (6e-04) b | 0.24 | 0.31 |

Abbreviations: $A_{1500}$ , photosynthesis at 1500 $\mu\text{mol m}^{-2}\text{s}^{-1}$ ( $\mu\text{mol m}^{-2}\text{s}^{-1}$ ); $\Phi$ , apparent quantum yield; LSP, light saturation point ( $\mu\text{mol m}^{-2}\text{s}^{-1}$ ); LCP, light compensation point ( $\mu\text{mol m}^{-2}\text{s}^{-1}$ ); $V_{cmax}$ , maximum carboxylation rate ( $\mu\text{mol m}^{-2}\text{s}^{-1}$ ); $J_{max}$ , maximum electron transport ( $\mu\text{mol m}^{-2}\text{s}^{-1}$ ); $R_D$ , respiration in the dark ( $\mu\text{mol m}^{-2}\text{s}^{-1}$ ); $R_L$ , respiration in the light ( $\mu\text{mol m}^{-2}\text{s}^{-1}$ ); $C_{area}$ , carbon per leaf area ( $\text{mg C cm}^{-2}$ ); $N_{area}$ , nitrogen per leaf area ( $\text{mg N cm}^{-2}$ ); C:N, carbon to nitrogen ratio; SLA, specific leaf area ( $\text{m}^{-2}\text{g}$ ); $Chl_{area}$ , total chlorophyll on an area basis ( $\mu\text{g cm}^{-2}$ );
Chl $a_{\text{area}}$ , area-based chlorophyll $a$ content ( $\mu\text{g cm}^{-2}$ ); Chl $b_{\text{area}}$ , area-based chlorophyll $b$ content ( $\mu\text{g cm}^{-2}$ ); Car $_{\text{area}}$ , area-based carotenoid content ( $\mu\text{g cm}^{-2}$ ); Chl:Car $_{\text{area}}$ , ratio of total chlorophyll content to carotenoids; Chl $a$ :Chl $b_{\text{area}}$ , ratio of chlorophyll $a$ to chlorophyll $b$ ; PRI, photochemical reflectance index.

#### Photosynthesis and Respiration

A significant positive relationship was observed between average PPFD and *A*_1500_ at BRF with high canopy foliage having a higher *A*_1500_ than low canopy foliage. No significant relationship was observed between *A*_1500_ and average PPFD at the FTE, nor were there any significant differences between the canopy positions (Fig 2a, Tables 2, S1 & S2). Across all samples at each location, mean *A*_1500_ was found to be 40% higher at BRF than at the FTE (10.8±0.5 at BRF vs. 7.7±0.4 µmol m^-2^ s^-1^ at the FTE; Table S2). The differences between BRF and the FTE in the relationship of *A*_1500_ to PPFD correspond to differences in several other parameters commonly extracted from light response curves. Namely the following were observed at BRF: a significant negative relationship between average PPFD and Φ (Fig 2b, Table 2), with significantly lower Φ in the high than the low canopy (Fig 2b, Tables S1 & S2); and significant positive relationships between average PPFD and LSP, LCP, *R*_D_, and *R*_L_ with higher values in the high than the low canopies (Figs 2c, 2d, 2e & 2f, Tables 2, S1 & S2). At the FTE, as in the case of *A*_1500_, no significant relationship was found between average PPFD and Φ, LCP, *R*_D_ or *R*_L_ (Figs 2b, 2d, 2e & 2f, Table 2); however, a significant positive relationship was found between average PPFD and LSP with a correspondingly higher LSP in the high canopy than the low canopy foliage at both locations (Fig 2c, Tables 2, S1 & S2). Across all samples at each location, Φ was significantly lower at the FTE than at BRF (0.03±0.001 at the FTE vs. 0.05±0.002 at BRF; Tables S1 & S2), and LSP, LCP, *R*_D_ and *R*_L_ were significantly higher at the FTE than at BRF (LSP = 644±14 at the FTE vs. 522±18.6 µmol m^-2^ s^-1^ at BRF; LCP = 90.7±4.5 at the FTE vs. 36.8±2.1 µmol m^-2^ s^-1^ at BRF (LCP means are backtransformed); *R*_D_ = 2.46±0.11 at the FTE vs. 1.98±0.14 µmol m^-2^ s^-1^ at BRF; *R*_L_ = 2.22±0.12 at the FTE vs. 1.58±0.16 µmol m^-2^ s^-1^ at BRF; Tables S1 & S2). In particular, the differences in mean *R*_L_ between the two locations were dramatic, with the FTE having 41% higher *R*_L_ than at BRF.

**Figure 2.**
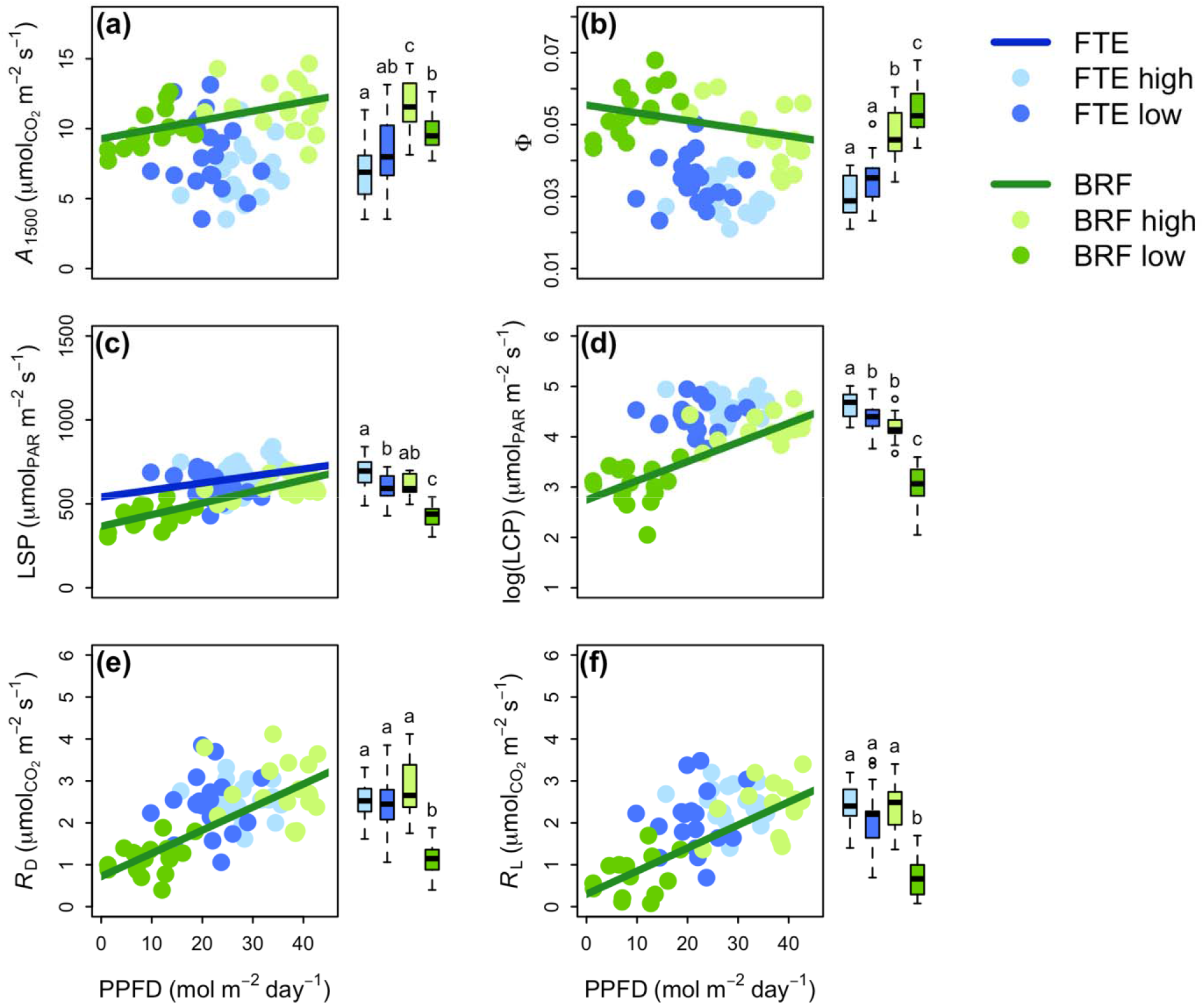
Linear relationships between average daily photosynthetic photon flux density (PPFD) over the measurement campaign and area-based foliar photosynthetic characteristics of white spruce from the Forest Tundra Ecotone (FTE; blue), Alaska, and Black Rock Forest (BRF; green), New York including photosynthesis at 1500 µmol m^-2^s^-1^ (*A*_1500_; 2a; FTE n = 38; BRF n = 33), apparent quantum yield (Φ; 2b; FTE n = 38; BRF n = 33), light saturation point (LSP; 2c; FTE n = 38; BRF n = 33), light compensation point (LCP; 2d; FTE n = 38; BRF n = 33), respiration in the dark (*R*_D_; 2e; FTE n = 38; BRF n = 36) and respiration in the light (*R*_L_; 2f; FTE n = 37; BRF n = 29). Light and dark colors at each location (BRF or FTE) represent high and low canopy positions, respectively. Parameter estimates of the linear mixed effects regression models, and statistical differences between slopes and intercepts are presented in Tables 2, S1 & S2. Regression lines are only shown for significant relationships (slope P<0.05). Also included are boxplots by canopy position and location for each parameter. Different letters represent significant differences between locations and canopy positions (P<0.05; Tables S1 & S2).

The parameters extracted from the *A-C*_i_ curves (*V*_cmax_ and *J*_max_ normalized to 25°C) also followed the positive relationship between PPFD and *A*_1500_ at BRF. Both *V*_cmax_ and *J*_max_ had positive relationships with PPFD and significantly higher values in the high than the low canopy at BRF (Fig 3, Tables 2, S1 & S2). However, the expected lack of a relationship with PPFD at the FTE was not observed. Instead, FTE *V*_cmax_ and *J*_max_ both had significant negative relationships with PPFD, with significantly lower values in the high than the low canopy. Consequently, when values from canopy positions were pooled to compare the two locations, no significant differences were observed between the FTE and BRF (*V*_cmax_ = 48.5±1.7 at the FTE and 50.3±2.2 µmol m^-2^ s^-1^ at BRF; *J*_max_ = 80.8±2.7 at the FTE and 85.9±3.4 µmol m^-2^ s^-1^ at BRF; Tables S1 & S2).

**Figure 3.**
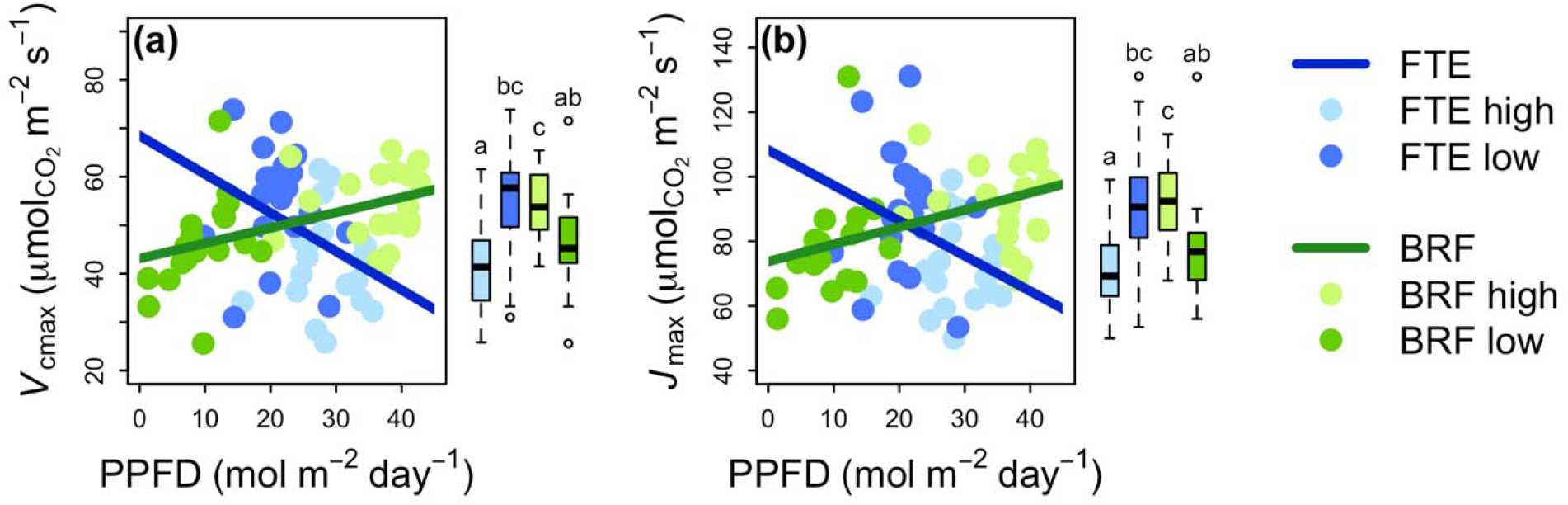
Linear relationships between average daily photosynthetic photon flux density (PPFD) and foliar respiratory characteristics on an area basis of white spruce from the Forest Tundra Ecotone (FTE; blue), Alaska, and Black Rock Forest (BRF; green), New York including the maximum rate of carboxylation (*V*_cmax_; 3a; FTE n = 39; BRF n = 37), and the maximum electron transport rate (*J*_max_; 3b; FTE n = 39; BRF n = 37). Light and dark colors at each location (BRF or FTE) represent high and low canopy positions, respectively. Parameter estimates of the linear mixed effects regression models, and statistical differences between slopes and intercepts are presented in Tables 2, S1, S2. Regression lines are only shown for significant relationships (slope P<0.05). Also included are boxplots by canopy position and location for each respiratory parameter. Different letters represent significant differences between locations and canopy positions (P<0.05; Tables S1 & S2).

#### Leaf traits

A significant negative relationship was observed between SLA and PPFD at BRF while no relationship was observed between SLA and PPFD at the FTE (Fig 4a, Table 2). Across canopy positions, high canopy foliage had significantly lower SLA at both locations (Fig 4a, Tables S1 & S2). The FTE had significantly lower SLA than BRF (32.0±1.1 at the FTE vs. 50.4±1.2 cm^2^g^-1^ at BRF, Tables S1 & S2). %N showed no significant relationships with PPFD, and no significant differences between canopy positions in either location; although the FTE did have a small but significantly lower %N than BRF (0.98±0.03 at the FTE vs. 1.1±0.04 at BRF; Fig 4c, Tables 2, S1 & S2). Similarly, C:N showed no significant relationships with PPFD and no significant differences between canopy positions in either location; however, C:N was significantly lower at BRF compared to the FTE (Tables 2, S1 & S2). Due to the strong relationship between SLA and PPFD, N_area_ also had a significant positive relationship with PPFD at BRF and no relationship at the FTE (Fig 4e, Table 2).

**Figure 4.**
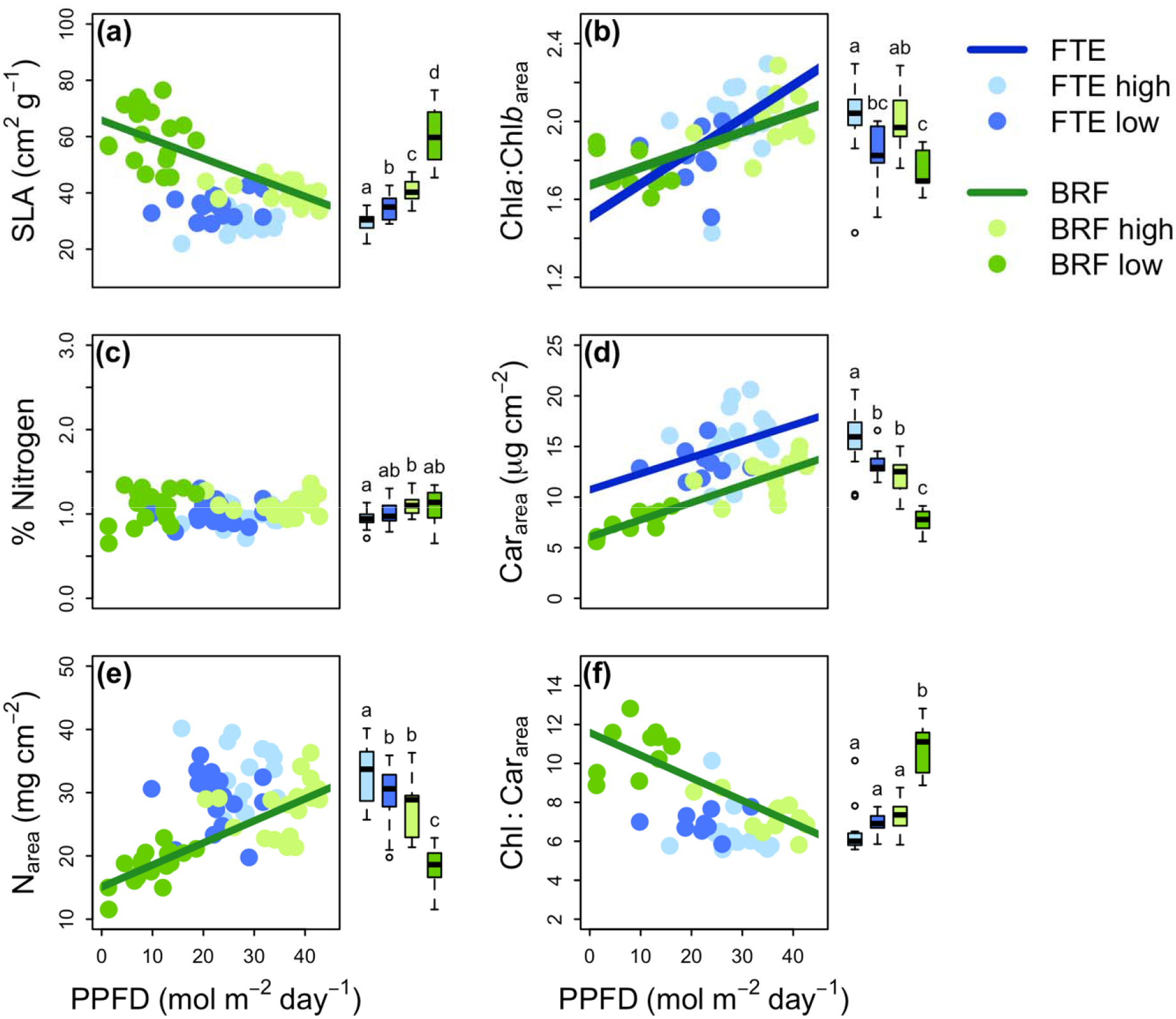
Linear relationships between average daily photosynthetic photon flux density (PPFD) and leaf traits of white spruce from the Forest Tundra Ecotone (FTE; blue), Alaska, and Black Rock Forest (BRF; green), New York including specific leaf area (SLA; 4a; FTE n = 37; BRF n = 38), the ratio of chlorophyll *a* to chlorophyll *b* (Chl*a*:Chl*b*_area_; 4b; FTE n = 25; BRF n = 22), % nitrogen (4c; FTE n = 37; BRF n = 38), carotenoids (Car_area_; 4d; FTE n = 25; BRF n = 22), nitrogen per leaf area (N_area_; 4e; FTE n = 37; BRF n = 38), and the ratio of chlorophylls *a* + *b* to carotenoids (Chl:Car_area_; 4c; FTE n = 25; BRF n = 22). Light and dark colors at each location (BRF or FTE) represent high and low canopy positions, respectively. Parameter estimates of the linear mixed effects models, and statistical differences between slopes and intercepts are presented in Tables 2, S1 & S2. Regression lines are only shown for significant relationships (slope P<0.05). Also included are boxplots by canopy position and location for each leaf trait. Different letters represent significant differences between locations and canopy positions (P<0.05; Tables S1 & S2).

#### Pigments and Spectral Indices

With regard to pigment concentrations, positive relationships were seen both at BRF and the FTE in the ratio of chlorophyll *a* and chlorophyll *b* (Chl*a*:Chl*b*_area_) and in the Carotenoids (Car_area_) with higher Chl*a*:Chl*b*_area_ and higher Car_area_ in the high than the low canopies (Figs 4b & 4d, Tables 2, S1 & S2). The expected decrease in the ratio of chlorophyll to carotenoids (Chl:Car_area_) with increasing PPFD was observed at BRF, but no relationship was found at the FTE (Fig 2f, Table 2). However, the FTE did have 46% more carotenoids than BRF (14.5±0.5 at the FTE vs. 9.9±0.8 µg cm^-2^ at BRF; Tables S1 & S2) and a significantly lower Chl:Car_area_ (6.7±0.2 at the FTE vs. 9.0±0.3 at BRF; Tables S1 & S2). Finally, PRI had the expected positive relationship with Chl:Car_area_ at BRF (Fig 5, Tables 3, S3) and it decreased significantly with increasing PPFD at BRF (Fig 5, Table 2). However, no relationships between PRI and Chl:Car_area_ or PPFD were found at the FTE (Fig 5, Tables 2, 3, S3).

**Figure 5.**
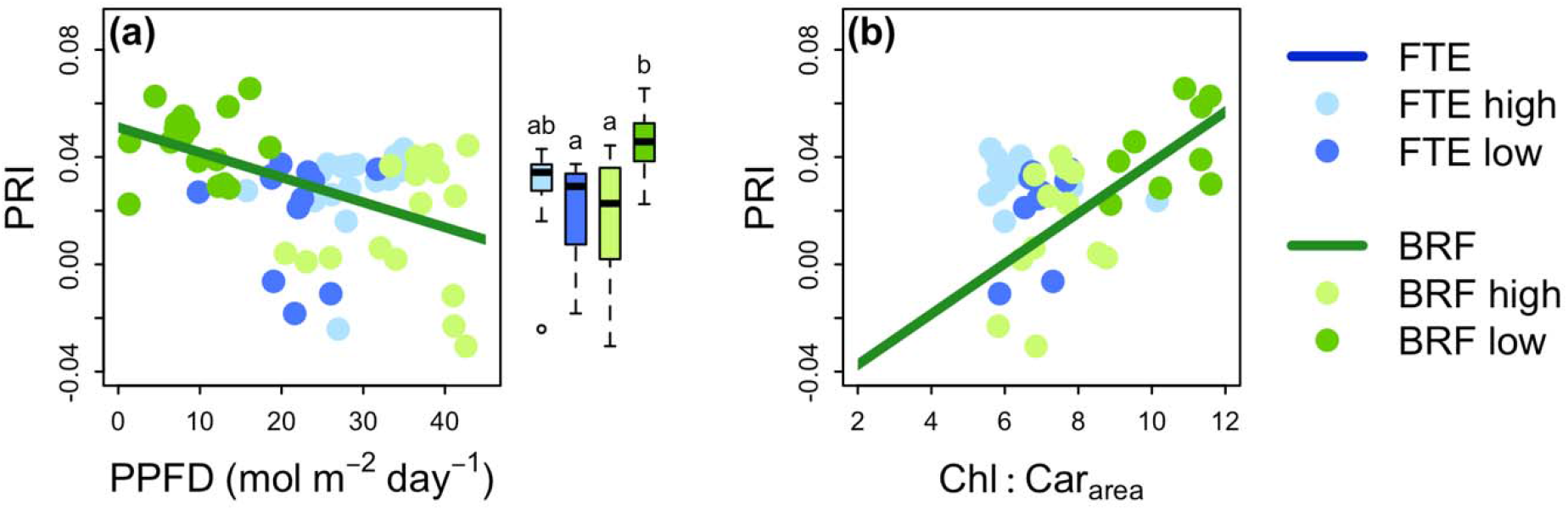
Linear relationships between photochemical reflectance index (PRI) and either average daily photosynthetic photon flux density (PPFD) (5a; FTE n = 30; BRF n = 34) or the ratio of chlorophylls *a* + *b* to carotenoids (Chl:Car_area_; 5b) of white spruce from the Forest Tundra Ecotone (FTE; blue), Alaska, and Black Rock Forest (BRF; green), New York. Light and dark colors at each location (BRF or FTE) represent high and low canopy positions, respectively. Parameter estimates of the linear mixed effects regression models, and statistical differences between slopes and intercepts are presented in Tables 2, S1 & S2. Regression lines are only shown for significant relationships (slope P<0.05). Also included are boxplots by canopy position and location for PRI. Different letters represent significant differences between locations and canopy positions (P<0.05; Tables S1 & S2).

**Table 3.** Parameter estimates for the intercepts and slopes of linear mixed effects models with tree as a random effect examining relationships between photosynthesis at 1500 µmol m^-2^s^-1^ (*A*_1500_), maximum rate of carboxylation (*V*_cmax_), respiration in the dark (*R*_D_), respiration in the light (*R*_L_) or total pigment content (Chl+Car_area_) and nitrogen per leaf area (N_area_); as well as the relationship between photochemical reflectance index (PRI) and the chlorophyll to carotenoid ratio (Chl:Car_area_). In all models tree was included as a random effect. Values in parentheses are standard SEs. Bolded and starred values are those that are significantly different from zero (***P<0.001; **P<0.01; *P<0.05; ^+^P<0.1). Significant differences between locations are indicated by different letters (P<0.05). Marginal and conditional coefficients of determination (r_m_^2^ and r_m_^2^, respectively) are also reported.

| Model | INTERCEPTS | | SLOPES | | $r_{\text{m}}^2$ | $r_{\text{c}}^2$ |
| --- | --- | --- | --- | --- | --- | --- |
|  | BRF | FTE | BRF | FTE |  |  |
| $A_{1500} \sim N_{\text{area}}^* \text{location}$ | **6.181 (1.250)\*\*\* a** | **8.130 (2.220)\*\*\* a** | **0.200 (0.049)\*\*\* a** | -0.024 (0.071) b | 0.5 | 0.64 |
| $V_{\text{cmax}} \sim N_{\text{area}}^* \text{location}$ | **28.828 (6.253)\*\*\* a** | **61.318 (12.106)\*\*\* b** | **0.934 (0.230)\*\*\* a** | -0.440 (0.383) b | 0.15 | 0.55 |
| $R_{\text{D}} \sim N_{\text{area}}^* \text{location}$ | -0.761 (0.410)<sup>+</sup> a | 0.714 (0.620) a | **0.118 (0.017)\*\*\* a** | **0.055 (0.020)\*\* b** | 0.49 | 0.49 |
| $R_{\text{L}} \sim N_{\text{area}}^* \text{location}$ | -0.908 (0.518)<sup>+</sup> a | 0.386 (0.723) a | **0.113 (0.021)\*\*\* a** | **0.059 (0.023)\* a** | 0.42 | 0.42 |
| $(\text{Chl}+\text{Car}_{\text{area}}) \sim N_{\text{area}}^* \text{location}$ | **62.309 (11.765)\*\*\* a** | **63.348 (22.613)\*\* a** | **1.436 (0.501)\*\* a** | **1.518 (0.712)\* a** | 0.39 | 0.52 |
| $\text{PRI} \sim \text{Chl}:\text{Car}_{\text{area}}^* \text{location}$ | **-0.060 (0.015)\*\*\* a** | 0.033 (0.023) b | **0.010 (0.002)\*\*\* a** | -0.0008 (0.003) b | 0.43 | 0.54 |

#### Relationships between gas exchange traits and nitrogen

As with PPFD, positive relationships were found between N_area_ and *A*_1500_, and N_area_ and *V*_cmax_ at BRF while no relationships were observed at the FTE (Fig 6a & 6b, Tables 3, S3). In contrast, both BRF and the FTE had positive relationships between N_area_ and *R*_D_ (Fig 6c, Tables 3, S3), between N_area_ and *R*_L_ (Tables 3, S3) and between N_area_ and total pigments (chl*a* + chl*b* + carotenoids; Fig 6d, Tables 3, S3).

**Figure 6.**
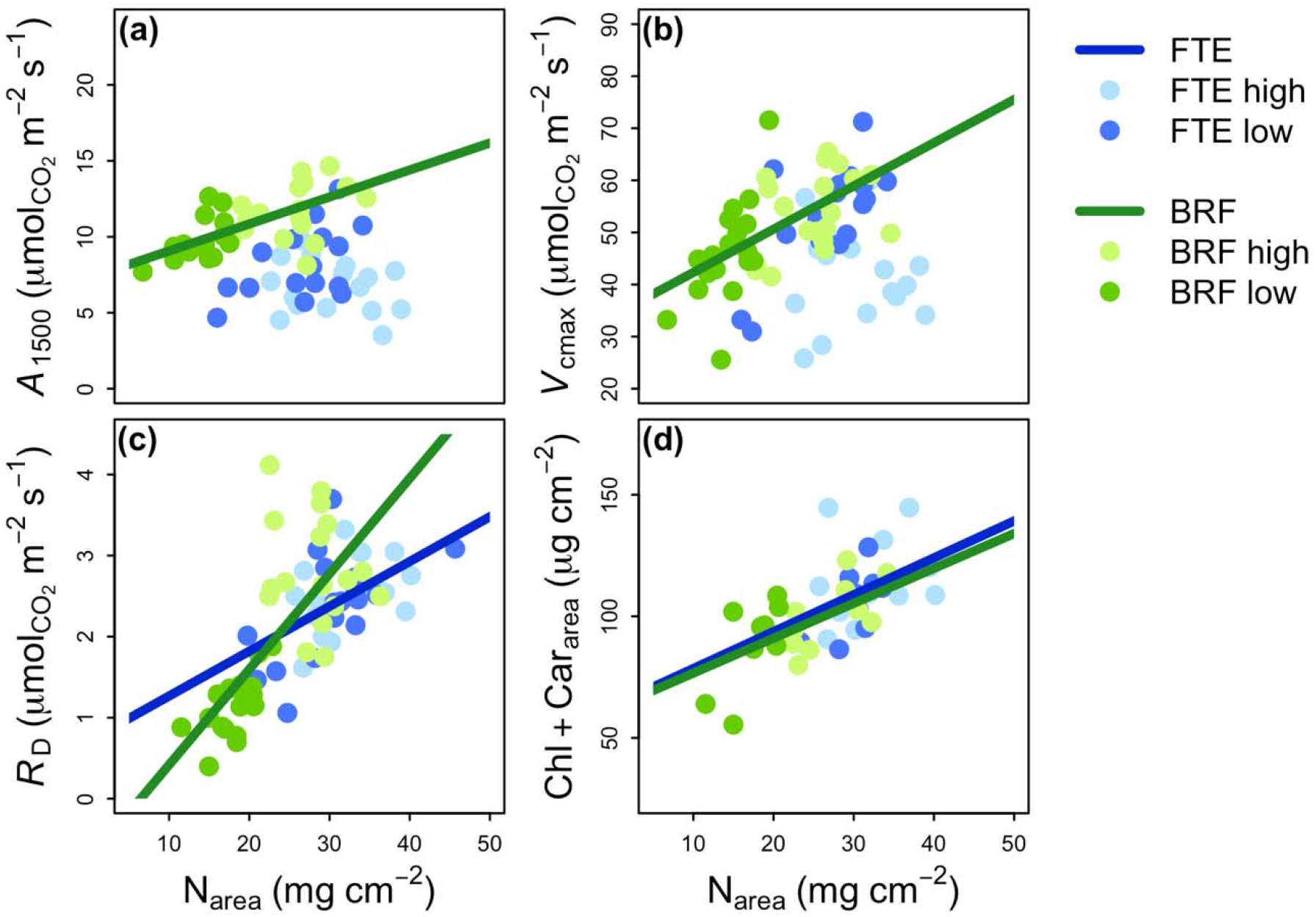
Linear relationships between photosynthesis at 1500 µmol m^-2^s^-1^ (*A*_1500_; 6a), maximum rate of carboxylation (*V*_cmax_; 6b), respiration in the dark (*R*_D_; 6c) and total pigment content (Chl+Car_area_; 6d) versus nitrogen per leaf area (N_area_) of white spruce from the Forest Tundra Ecotone (FTE; blue), Alaska and Black Rock Forest (BRF; green), New York. Light and dark colors at each location (BRF or FTE) represent high and low canopy positions, respectively. Parameter estimates of the linear mixed effects regression models, and statistical differences between slopes and intercepts are presented in Tables 3 & S3. Regression lines are only shown for significant relationships (slope P<0.05).

### Carbon balance at the range extremes of white spruce

The collection of gas exchange parameters such as *V*_cmax_ and *J*_max_, which are commonly used to predict photosynthesis, together with values of respiration at 25°C (*R*_25_) previously reported by Griffin *et al*. (2022), provide a unique opportunity to examine the carbon balance across the range extremes of white spruce. We find that, when all parameters are normalized to 25°C, the ratios of *V*_cmax_ and *J*_max_ to *R*_25_ are 228% higher at BRF than at the FTE (Fig 7, Tables S1 & S2; *V*_cmax_/*R*_25_ = 21.4±1.39 at the FTE vs. 70.2±7.47 at BRF; *J*_max_/*R*_25_ = 36.5±2.45 at the FTE vs. 119.9±13.21 at BRF). No significant differences were observed in the ratios between canopy positions at each location, although there was a trend for larger ratios in the low canopy than the high canopy at BRF (Fig 7, Tables S1 & S2).

**Figure 7.**
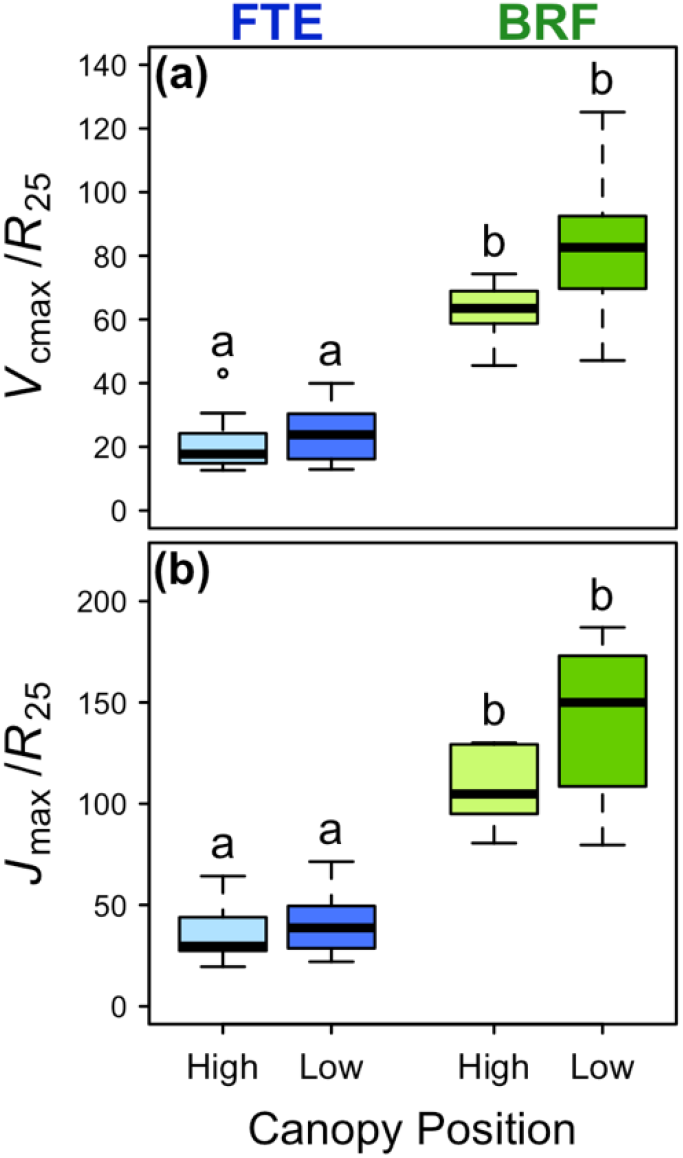
Ratios of (a) maximum rate of carboxylation to respiration at 25°C from Griffin *et al*. (2022) (*V*_cmax_/*R*_25_) and (b) maximum electron transport rate to respiration at 25°C from Griffin *et al*. (2022) (*J*_max_/*R*_25_) of white spruce from high and low canopy positions at the Forest Tundra Ecotone (FTE, blue), Alaska, and Black Rock Forest (BRF, green), New York. Boxplots show the median and first and third quartiles. Whiskers display the range of groups with individual points representing outliers falling outside 1.5 times the interquartile range. Different letters represent significant differences between locations and canopy positions (P<0.05; Tables S1 & S2).

## DISCUSSION

### Local environmental conditions drive differences in tree structure between white spruce range extremes

Vertical canopy gradients of foliar physiological and biochemical traits differ between white spruce growing at its northern- and southern-most range limits. Whereas steep canopy gradients in physiological functioning are found at the southern range limit in Black Rock Forest (BRF), a clear lack of variation in foliar physiological capacity is observed between the high and low canopies of white spruce at its northernmost range extreme in the Forest-Tundra Ecotone (FTE). As our two study locations delimit the distributional range of white spruce and are separated by a distance of more than 5000 km, these differences are likely in response to local environmental conditions. Solar zenith angles, day length, air and soil temperatures, soil moisture and vapor pressure deficit vary substantially between these two locations. Of these environmental conditions, the impacts of local light environment are of particular interest for the following reasons: 1) light energy is a critical driver of photosynthetic carbon gain (Hirose & Werger 1987); 2) abundant evidence shows that canopy gradients in photosynthetic traits are strongly correlated to gradients in irradiance throughout tree crowns (Field 1983; Hirose & Werger 1987; Ellsworth & Reich 1993; Bond *et al*. 1999; Niinemets, Keenan & Hallik 2015), but little work assesses the impacts of latitudinal location on these gradients within a single species; and 3) these gradients are key parameters in ecosystem models used to calculate gross primary productivity across latitudinal space (Bonan *et al*. 2012; Rogers *et al*. 2017). Day length and solar zenith angles increase and light intensity decreases as one moves towards the northernmost latitudes of the globe (Nilsen 1983; Slaughter & Viereck 1986). These local light environments are further modified by differences in forest and tree crown structure between the two locations. Trees at the southern location exist within a complex forest canopy comprised of deciduous hardwood species where they must compete for light energy and other above-ground resources. Consequently white spruce tree crowns are distinctly conical in shape with a wide dripline area in the southern location, presumably to minimize self-shading and maximize light energy capture. In contrast, trees at the northern location are widely spaced with narrow and untapering tree crowns. The open forest canopy minimizes competition for resources such as light. Furthermore, the cylindrical shape promotes more equal exposure of foliage to irradiance in a region of the globe with extremely high solar zenith angles, ultimately increasing the efficiency of light capture for photosynthesis (Kuuluvainen 1992). Overall, these solar and structural differences lead to a narrowing of the intra-canopy light gradients as one moves from the southern (where light availability is ∼35 and ∼9 mol m^-2^ day^-1^ at the high and low canopies, respectively) to the northern (where light availability is ∼29 and ∼21 mol m^-2^ day^-1^ at the high and low canopies, respectively) latitudinal limits of this species.

### Vertical gradients in physiological function are coordinated to latitudinal differences in light environment

Inherent to the structural differences of trees in these two locations is the need for tree crowns to organize their physiological and biochemical traits in order to optimize photosynthetic carbon gain (Hirose & Werger 1987; Niinemets *et al*. 1998). Therefore, at BRF, where there are strong intra-canopy light gradients, we find strongly positive relationships between PPFD and *A*_1500_, *J*_max_ and *V*_cmax_ indicating the upregulation of the light reactions (in which RuBP is produced) and the carbon reactions (in which rubisco acts on RuBP to fix carbon) of photosynthesis in the high canopy (Givnish 1988; Valladares & Niinemets 2008). The elevated metabolic activity in the high canopy results in higher respiratory activity in order to support the costs of growth and protein maintenance (Reich *et al*. 1998; Lewis, McKane, Tingey & Beedlow P. A. 2000; Griffin, Tissue, Turnbull, Schuster & Whitehead 2001). Biochemical traits also follow the inherent light gradients. High light needles have lower SLA and a greater N_area_ reflecting the greater leaf thickness of high light leaves from added palisade layers and the presence of nitrogen in both the enzyme rubisco and in chlorophyll (Evans 1989; Smith, Vogelmann, DeLucia, Bell & Shepherd 1997; Evans & Poorter 2001). Finally, high light leaves have greater Car_area_ and a smaller Chl:Car_area_ ratio reflecting the need for greater photoprotection from excess light energy (Demmig-Adams & Adams III 1992). In all cases, biochemical and physiological traits of low latitude white spruce conform to the established literature showing strong correlations with canopy gradients of irradiance. At the high latitude location where intra-canopy light gradients become less pronounced, we see the absence of canopy gradients in many physiological and biochemical traits. What is most striking is the lack of relationships between irradiance and each of photosynthesis (*A*_1500_), light respiration, and dark respiration. Together with the lack of relationships between irradiance and the apparent quantum yield, the light compensation point, specific leaf area and N_area_, these findings suggest that the uneven distribution of metabolic activity throughout the canopy required to optimize photosynthetic carbon gain in correspondence with canopy irradiance gradients at low latitudes does not occur at high latitudes where irradiance varies minimally throughout the canopy profile.

The parameters discussed above suggest that northern latitude trees have few distinguishable canopy gradients in photosynthetic physiology. However, positive relationships between irradiance and the ratio of Chl*a* to Chl*b*, the light saturation point and carotenoid concentrations suggest that even though the magnitude of the light gradient narrows at northern latitudes, there is still a subtle intra-canopy light gradient to which pigments are responding. Thus, in the higher light upper canopy we see greater investments in Chl*a* than Chl*b* to maximize light capture and a greater investment in carotenoids to dissipate excess light energy and protect photosynthetic machinery (Dale & Causton 1992; Hikosaka & Terashima 1995; Kitajima & Hogan 2003; Ruban 2015; Magney *et al*. 2016; Scartazza *et al*. 2016). It is possible that these subtle canopy gradients in pigments may be a result of the 24 hours of light exposure during the growing season at the FTE.

In contrast to the lack of relationships found between irradiance and the majority of traits, and the subtle positive relationships found between irradiance and pigments at treeline, we find a surprising downregulation of *J*_max_ and *V*_cmax_ in the high canopy foliage at the FTE. It is unclear why we find these negative relationships between irradiance and the component processes of photosynthesis at the northern locations; however, there are several possible explanations and hypotheses that could be further explored. One possibility is that the combination of diminutive vertical canopy gradients in light, SLA and nitrogen, the latter of which is critical to the performance of *J*_max_ and *V*_cmax_ because of the presence of nitrogen in chlorophyll and the enzyme rubisco (Evans 1989; Smith *et al*. 1997; Evans & Poorter 2001), has led to the observed negative relationships. However, strong relationships do exist between nitrogen and total pigment concentrations even though there is no apparent relationship between nitrogen and *V*_cmax_. Furthermore, we still find weak positive relationships between irradiance and both carotenoid concentrations and Chl*a*:Chl*b* even with a narrow light gradient. Consequently, we conclude that a more mechanistic explanation must exist for the puzzling downregulation of *V*_cmax_ and *J*_max_. While in this study we focused on the latitudinal relationships between canopy light environments and foliar physiological traits, it is possible that intra-canopy gradients in other environmental conditions may also impact *V*_cmax_ and *J*_max_ measurements (Martin, Hinckley, Meinzer & Sprugel 1999; Zweifel, Böhm & Häsler 2002; Bauerle, Bowden, Wang & Shahba 2009; Bachofen, D’Odorico & Buchmann 2020; Buckley 2021). Perhaps of particular relevance with respect to rubisco carboxylation is the reliance of this process on the enzyme rubisco, whose activity is well-known to be temperature dependent (Yamori & von Caemmerer 2009; Benomar *et al*. 2018). Work at alpine treelines has clearly documented that as trees grow in stature, their foliage becomes increasingly coupled to atmospheric conditions such as temperature and wind stress (Körner 2012). These stresses may be detrimental to high canopy photosynthetic performance (lowering *V*_cmax_ and *J*_max_). In contrast, low canopy foliage is less coupled to atmospheric conditions and may therefore be more protected from temperature and wind extremes (Martin *et al*. 1999; Germino & Smith 1999; Johnson, Germino & Smith 2004; Maguire *et al*. 2019).

Temperature and wind stress are not the only environmental conditions likely to cause reduced gradients in physiological functioning. Hydraulic conductance has also been found to vary in response to water availability, impacting canopy gradients in nitrogen and photosynthesis (Peltoniemi *et al*. 2012). It is equally possible that winter temperature extremes may impact hydraulic function in exposed foliage. Consequently, throughout the canopy of Alaskan trees there may be a constant balancing act with regard to the resource allocation necessary to improve whole canopy photosynthetic carbon gain. Two factors compete with each other: namely the need to devote resources to light capture in the high canopy where irradiance is slightly more available; and the need to devote resources either to foliage protection in the high canopy or increased carbon gain in the low canopy when harsh conditions prevail. The result, ultimately, is what we find across the majority of physiological and biochemical traits, i.e., a lack of variation in photosynthetic carbon gain across the high and low canopy in high latitude trees. Without detailed information on temperature, wind, and hydraulic profiles throughout the canopy, this hypothesis cannot be fully tested. Yet, it seems clear from the linkages between irradiance, photosynthesis and tree structure across the latitudinal extremes that the environment is an integral driver of species’ geographic range. We suggest that expanding to examine additional environmental conditions such as temperature, wind and water stress may be a compelling future direction, thereby adding to our overall understanding of the mechanisms controlling canopy carbon exchange across the massive forest tundra ecotone and more broadly across latitudinal gradients.

The accurate portrayal of canopy gradients is important to the precise prediction of carbon exchange and gross primary productivity (GPP) (Sands 1995; Bonan *et al*. 2012). However, the lack of variation in photosynthesis at the FTE suggests that complex canopy models used to account for canopy self-shading at mid-latitudes may not be necessary to model GPP in trees from high latitudes. The lack of variation in the photochemical reflectance index (PRI) across vertical canopy positions at the FTE similarly suggests that accounting for canopy self-shading may be less critical at northern latitudes as opposed to mid-latitudes. PRI has previously been demonstrated to be an accurate predictor of whole canopy gross primary productivity in broad-leaved species when used in concert with terrestrial Light Detection and Ranging (LiDAR) to account for canopy self-shading (Hilker *et al*. 2008; Coops, Hermosilla, Hilker & Black 2017). Although our own study does not employ LiDAR, the presence of canopy differences in PRI at the southernmost location suggests that such a technique could work equally well in mid-latitude coniferous forests. Furthermore, the fact that PRI mirrors *A*_1500_ regardless of latitude suggests that this remotely sensed index may be a valuable future tool for accurately assessing gross primary productivity of other species with different structural properties and with large latitudinal distributions.

### Physiological and biochemical traits may contribute to the range limits of white spruce

The detailed data on the physiology and biochemistry of white spruce not only allow for the examination of canopy differences across latitudinal range extremes, but also for the examination of locational differences in traits that may further illuminate possible drivers contributing to the range limits of this important species. Here, we observed important differences in physical and biochemical traits such as SLA, % N, C:N and carotenoid concentration between the southern and northern range extremes. Most relevant to the above discussion of light environment and its impacts on photosynthetic physiology is the finding of significantly higher carotenoid concentrations in needles of northern white spruce compared to southern white spruce. This indicates a greater investment in photoprotection at northern latitudes during the growing season. Elevated carotenoids at the northern range extreme may be in response to the 24 hour photoperiod at the Arctic tundra boreal forest ecotone. Past studies have found that very high levels of light protecting xanthophyll pigments can be a feature of Arctic plants subjected to continuous light (Magney *et al*. 2019). Lower overall SLA in Alaskan needles compared to BRF needles indicates that foliage at the northern edge of the distribution is thicker or denser. Although decreasing SLA is commonly found in response to increasing light availability due to the added density of additional photosynthetic machinery (as seen throughout the BRF canopy), SLA is also commonly positively correlated to ambient growth temperatures (Atkin, Loveys, Atkinson & Pons 2006; Poorter, Niinemets, Poorter, Wright & Villar 2009). Thus, a low SLA at the FTE may indicate a greater investment in needle structure rather than photosynthetic machinery, a hypothesis that is corroborated by our finding of higher carbon to nitrogen ratios in Alaskan needles (González-Zurdo, Escudero, Babiano, García-Ciudad & Mediavilla 2016). Investment in structure is potentially adaptive in the harsh forest tundra ecotone where needles must survive multiple years of winter temperatures, snow or ice accumulation and windblown ice abrasion. Leaf nitrogen was also found to be slightly but significantly lower in needles from the northern range limit. This result confirms the findings of Griffin *et al*. (2022) and agrees with past studies that show negative correlations between nitrogen and both temperature and latitude (Körner 1989; Yin 1993; Reich & Oleksyn 2004). Nitrogen biogeochemistry is intimately linked to environmental and ecological factors. For example, harsh temperatures and unique hydrologic and permafrost dynamics inherent to the Arctic can slow decomposition rates and depress nitrogen fixation (Schimel & Stuart Chapin 1996; Schimel, Bilbrough & Welker 2004). The result is the limited availability of nitrogen in the harsh northern latitudes of the globe. The differences in biochemical and structural traits between our two study locations hint at environmental drivers, in addition to light availability, that should be considered in future studies examining latitudinal differences in canopy gradients.

Finally, in addition to biochemical differences between locations, metabolic differences are key in understanding the mechanisms controlling the range limits of this species. Previous work in white spruce shows this species to have extremely high rates of dark respiration at its northern compared to southern range limit, a finding that has led to the conclusion that respiratory carbon loss may be a crucial factor constraining the northern range limit of this species (Griffin *et al*. 2022). We find similar patterns in the current study, with significantly higher rates of dark respiration (as estimated from the light response curves) in white spruce in Alaska than in New York. Furthermore, we find that these differences are apparent not only in dark respiration, but also in light respiration. Given that the northern range extreme of this species experiences 24 hours of sunlight during the majority of the growing season, the greater rates of light respiration in FTE compared to BRF trees provides important corroboration of the constancy of the extreme respiratory cost at this northern range limit and the role it may play in determining the location of northern treeline. It is likely that these high respiratory fluxes to the atmosphere are related to high maintenance costs given the harsh environmental conditions of the region.

Northern white spruce are not only characterized by high carbon losses. The additional information on photosynthetic processes provided here further suggests that these carbon costs are not matched by similarly high carbon gains. We do not directly compare the ratios of photosynthetic carbon gain to respiratory carbon loss due to the use of different measurement temperatures chosen to reflect ambient conditions in each location, but the ability to normalize both *V*_cmax_ and *J*_max_ to 25°C allows us to assess the carbon balance of white spruce at its latitudinal range extremes. The results of this comparison are dramatic and demonstrate that northern trees function within a much narrower margin to maintain a positive carbon balance. It seems that it is not only high respiratory costs that result in slow growth rates of white spruce (Jensen et al., *pers. com*.) and constrain the latitudinal range limit of northern treeline. Instead, our study suggests that it is the combined effect of high respiration and low *V*_cmax_ and *J*_max_ that ultimately limit the northern range of this widely distributed and important boreal species.

### Conclusions

White spruce spans a massive distribution, the extremes of which are characterized by vastly different environmental conditions. Here we demonstrate that differences in local light environment correlate to differences in vertical canopy physiology, resource allocation and crown structure. Thus, at high latitudes where light intensity is low, solar zenith angles are high and day length is long, there is a clear lack of variation between high and low canopy white spruce traits. In contrast, white spruce at lower latitudes exist in a complex forest canopy of deciduous hardwood species and experience greater differences in local and canopy light environments leading to steep gradients in physiological and biochemical traits between high and low canopy positions. Consequently, accounting for self-shading may be less critical for predicting gross primary productivity at northern relative to southern latitudes. Our findings point to light environment as an important driver in latitudinal variation in physiology within a single species and emphasize the importance of the connection between environment and physiology in determining a species’ distributional range. Furthermore, our findings highlight that additional environmental drivers such as temperature, and water availability may also contribute both to canopy and latitudinal differences in physiological traits.

These data not only allowed us to examine the role of physiology in determining a species’ distributional range, but they also permitted us to question the role of a species’ carbon balance in determining the extreme limit of tree growth form. Previous work has highlighted that extreme respiratory costs may be a critical component leading to the latitudinal location of northern treeline. Here we demonstrate that it is not only extreme respiratory costs but also minimal photosynthetic carbon gains that together may determine the latitudinal location of the world’s largest ecotone and the transition from boreal forest to treeless tundra.

## ACKNOWLEDGEMENTS

This work was supported by NASA ABoVE grant NNX15AT86A and the Arctic LTER (NSF Grant No. 1637459 & 2220863). We thank Sarah Sacket from the NASA ABoVE support team in Fairbanks, AK, for her energetic and prompt logistical support during Alaska field campaigns. At Black Rock Forest, we also wish to acknowledge the unflagging support of BRF staff including Ben Brady and Matthew Munson as well as Dr. William Schuster, the Executive Director of Black Rock.

## AUTHOR CONTRIBUTION

SCS, KLG, NTB, LAV and JUHE designed the research. They were assisted in data collection by SGB, LM, and JJ. SCS analyzed the data and wrote the first draft of the manuscript. All authors edited and approved the final version for submission.

## DATA AVAILABILITY

Data will be archived with ORNL DAAC upon acceptance of the manuscript.

## SUPPLEMENTAL INFORMATION

**Methods S1** *Linear mixed effects analysis of the relationships between physiological variables and N*_*area*_ *or PRI and Chl:Car*_*area*_

The relationships between N_area_ and other physiological parameters by location presented in Table 3 and Figures 5 & 6 were assessed with a linear mixed effects model in which the interaction term allowed for the slope to vary by location (Black Rock Forest (BRF) or the Forest Tundra Ecotone (FTE)). Multiple measurements per tree were accounted for by including a random effect for each individual tree. The full model can be written as such:

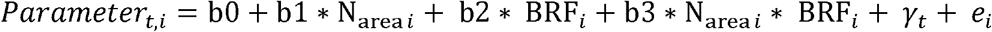

Where *i* represents each individual measurement taken, *t* represents each tree measured, BRF is a binary variable (e.g. BRF is 1 if the individual is from BRF, and 0 if it is not (i.e. from the FTE instead)), *γ* is a normally distributed random variable for different trees: *γ* ∼ N(0, *σ*), and *e* is the standard normally distributed random error term: *e* ∼ N(0, *σ*_*e*_).

This means that, for trees from the FTE, the equation is:

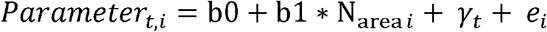

Whereas, for trees from BRF, the equation is:

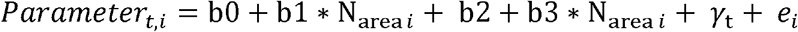

**Table S1.** Summaries of the mixed effects models in which parameters were modeled as a function of the interaction between canopy position and location with a random effect for each individual tree. P-values are estimated using the Kenward-Roger approximation of the degrees of freedom as suggested by (Halekoh & Højsgaard 2014). See table 2 for explanations of abbreviations and units. P<0.05 are bolded.

|  | Canopy |  | Location |  | Canopy x Location |  |
| --- | --- | --- | --- | --- | --- | --- |
|  | *F*-value | *p*-value | *F*-value | *p*-value | *F*-value | *p*-value |
| $A_{1500}$ | 0.14 | 0.7106 | 26.14 | **<0.001** | 13.28 | **<0.001** |
| $\Phi$ | 15.57 | **<0.001** | 60.94 | **<0.001** | 0.68 | 0.4129 |
| $LSP_{1500}$ | 59.20 | **<0.001** | 27.38 | **<0.001** | 11.31 | **0.0015** |
| $\log(LCP)$ | 94.83 | **<0.001** | 139.98 | **<0.001** | 41.65 | **<0.001** |
| $V_{cmax}$ | 1.35 | 0.2506 | 0.23 | 0.6381 | 23.37 | **<0.001** |
| $J_{max}$ | 0.65 | 0.4233 | 1.00 | 0.3333 | 23.82 | **<0.001** |
| $R_D$ | 55.17 | **<0.001** | 7.61 | **0.0146** | 46.38 | **<0.001** |
| $R_L$ | 49.13 | **<0.001** | 10.09 | **0.006** | 25.33 | **<0.001** |
| % Carbon | 1.55 | 0.2194 | 1.41 | 0.2482 | 12.79 | **<0.001** |
| % Nitrogen | 0.09 | 0.7625 | 5.97 | **0.0265** | 1.88 | 0.1757 |
| $C_{area}$ | 70.78 | **<0.001** | 148.02 | **<0.001** | 4.19 | **0.0452** |
| $N_{area}$ | 77.16 | **<0.001** | 25.95 | **<0.001** | 19.83 | **<0.001** |
| $\log(C:N)$ | 0.05 | 0.8181 | 6.25 | **0.0236** | 1.58 | 0.2146 |
| SLA | 90.17 | **<0.001** | 134.52 | **<0.001** | 31.35 | **<0.001** |
| $Chl_{area}$ | 2.68 | 0.1125 | 5.32 | **0.0363** | 0.02 | 0.8997 |
| $Chla_{area}$ | 8.12 | **0.0081** | 6.27 | **0.0246** | 0.05 | 0.825 |
| $Chlb_{area}$ | 0.10 | 0.7494 | 3.09 | 0.0997 | 0.0004 | 0.9843 |
| $Car_{area}$ | 49.82 | **<0.001** | 22.34 | **<0.001** | 3.56 | 0.0703 |
| $Chl:Car_{area}$ | 53.41 | **<0.001** | 31.14 | **<0.001** | 23.67 | **<0.001** |
| $Chla:Chlb_{area}$ | 34.49 | **<0.001** | 0.37 | 0.5532 | 0.42 | 0.5213 |
| PRI | 7.37 | **0.0094** | 0.33 | 0.5751 | 20.94 | **<0.001** |
| $V_{cmax}/R_{25}$ | 5.32 | **0.0319** | 91.24 | **<0.001** | 0.15 | 0.7041 |
| $J_{max}/R_{25}$ | 3.68 | 0.0693 | 85.21 | **<0.001** | 0.60 | 0.4484 |

**Table S2.** Means ± 1 standard error for physiological, leaf and spectral traits. Several traits report back-transformed means including the light compensation point (LCP) and the ratio of carbon to nitrogen (C:N). Significant differences are marked by different letters (P < 0.05) with differences between locations marked by capital letters and differences between location and canopy positions marked by lowercase letters. See table 2 for explanations of abbreviations and units.

|  | BRF | FTE | BRF low | BRF high | FTE low | FTE high |
| --- | --- | --- | --- | --- | --- | --- |
| $A_{1500}$ | 10.8 (0.5) A | 7.7 (0.4) B | 9.8 (0.6) b | 11.7 (0.6) a | 8.3 (0.5) bc | 7.0 (0.5) c |
| $\Phi$ | 0.05 (0.002) A | 0.03 (0.001) B | 0.05 (0.002) a | 0.05 (0.002) b | 0.03 (0.002) c | 0.03 (0.002) c |
| $LSP_{1500}$ | 522.0 (18.6) A | 643.9 (14.1) B | 427.4 (22.5) a | 616.7 (22.2) bc | 605.6 (17.8) b | 682.2 (18.4) c |
| LCP | 36.8 (2.1) A | 90.7 (4.5) B | 20.7 (1.6) a | 65.1 (5.0) b | 79.7 (5.3) b | 103.1 (7.2) c |
| $V_{cmax}$ | 50.3 (2.2) A | 48.5 (1.7) A | 46.2 (2.7) ab | 54.5 (2.7) c | 54.7 (2.2) bc | 42.3 (2.35) a |
| $J_{max}$ | 85.9 (3.4) A | 80.8 (2.7) A | 78.7 (4.1) ab | 93.1 (4.2) c | 90.1 (3.5) bc | 71.5 (3.7) a |
| $R_D$ | 1.98 (0.14) A | 2.46 (0.11) B | 1.14 (0.17) a | 2.82 (0.17) b | 2.41 (0.13) b | 2.5 (0.14) b |
| $R_L$ | 1.58 (0.16) A | 2.22 (0.12) B | 0.77 (0.19) a | 2.38 (0.19) b | 2.05 (0.14) b | 2.39 (0.14) b |
| % Carbon | 47.96 (0.50) A | 48.63 (0.30) A | 47.64 (0.51) a | 48.27 (0.51) b | 48.81 (0.32) ab | 48.45 (0.32) ab |
| % Nitrogen | 1.1 (0.04) A | 0.98 (0.03) B | 1.09 (0.05) ab | 1.12 (0.05) b | 1.00 (0.03) ab | 0.96 (0.04) a |
| $C_{area}$ | 1007.2 (33.5) A | 1547.8 (29.5) B | 813.4 (42.8) a | 1201.1 (42.1) b | 1430.0 (38.2) c | 1665.5 (40.9) d |
| $N_{area}$ | 23.1 (1.3) A | 31.2 (0.9) B | 18.3 (1.4) a | 27.9 (1.4) bc | 29.7 (1.0) b | 32.7 (1.1) c |
| C:N | 43.9 (1.8) A | 49.9 (1.5) B | 44.4 (2.0) ab | 43.3 (2.0) a | 48.9 (1.7) ab | 50.9 (1.8) b |
| SLA | 50.4 (1.2) A | 32.0 (1.1) B | 60.2 (1.5) d | 40.5 (1.5) c | 34.5 (1.4) b | 29.5 (1.5) a |
| $Chl_{area}$ | 84.6 (3.9) A | 96.0 (3.1) B | 81.4 (4.8) a | 87.7 (4.5) ab | 93.3 (4.6) ab | 98.7 (3.4) b |
| $Chla_{area}$ | 55.0 (2.5) A | 62.9 (2.0) B | 51.5 (3.0) a | 58.4 (2.9) ab | 60.0 (2.9) ab | 65.9 (2.2) b |
| $Chlb_{area}$ | 29.6 (1.6) A | 33.1 (1.3) A | 29.8 (1.9) a | 29.3 (1.8) a | 33.3 (1.9) a | 32.9 (1.4) a |
| $Car_{area}$ | 9.9 (0.8) A | 14.5 (0.5) B | 7.6 (0.9) a | 12.1 (0.8) b | 13.2 (0.7) b | 15.7 (0.6) c |
| $Chl:Car_{area}$ | 9.0 (0.3) A | 6.7 (0.2) B | 10.7 (0.4) b | 7.3 (0.3) a | 7.0 (0.3) a | 6.4 (0.3) a |
| $Chla:Chlb_{area}$ | 1.87 (0.05) A | 1.92 (0.04) A | 1.75 (0.06) a | 2.00 (0.05) bc | 1.81 (0.05) ab | 2.02 (0.04) c |
| PRI | 0.03 (0.004) A | 0.02 (0.004) A | 0.04 (0.005) b | 0.02 (0.005) a | 0.02 (0.005) a | 0.03 (0.004) ab |
| $V_{cmax}/R_{25}$ | 70.2 (7.5) A | 21.4 (1.4) B | 23.4 (1.88) a | 19.5 (1.67) a | 61.7 (8.35) b | 79.9 (10.81) b |
| $J_{max}/R_{25}$ | 119.9 (13.2) A | 36.5 (2.4) B | 38.7 (3.06) a | 34.5 (2.98) a | 105.8 (14.08) b | 135.8 (18.07) b |

**Table S3.** Fixed effects parameter estimates and their significance for the linear mixed effects models assessing relationships between physiological variables and N_area_ or PRI and Chl:Car_area_ with an interaction term for location and a random effect for individual tree (taking into account multiple measurements taken on each tree at the high and low canopy positions). The full models can be seen in the Methods S1 available in the supplemental information.

| Model | b0 | b1 | b2 | b3 |
| --- | --- | --- | --- | --- |
| $A_{1500} \sim N_{\text{area}} * \text{location}$ | **0.0007** | 0.7342 | 0.4477 | **0.0118** |
| $V_{\text{cmax}} \sim N_{\text{area}} * \text{location}$ | **0.00001** | 0.2553 | **0.0207** | **0.0031** |
| $R_D \sim N_{\text{area}} * \text{location}$ | 0.2539 | **0.0060** | 0.0516 | **0.0190** |
| $R_L \sim N_{\text{area}} * \text{location}$ | 0.5956 | **0.0131** | 0.1514 | 0.0921 |
| $(\text{Chl} + \text{Car}_{\text{area}}) \sim N_{\text{area}} * \text{location}$ | **0.0080** | **0.0395** | 0.9677 | 0.9253 |
| $\text{PRI} \sim \text{Chl:Car}_{\text{area}} * \text{location}$ | 0.1590 | 0.8108 | **0.0016** | **0.0073** |

